# Hippocampal replay reflects specific past experiences rather than a plan for subsequent choice

**DOI:** 10.1101/2021.03.09.434621

**Authors:** Anna K. Gillespie, Daniela A. Astudillo Maya, Eric L. Denovellis, Daniel F. Liu, David B. Kastner, Michael E. Coulter, Demetris K. Roumis, Uri T. Eden, Loren M. Frank

## Abstract

Executing memory-guided behavior requires both the storage of information about experience and the later recall of that information to inform choices. Awake hippocampal replay, when hippocampal neural ensembles briefly reactivate a representation related to prior experience, has been proposed to critically contribute to these memory-related processes. However, it remains unclear whether awake replay contributes to memory function by promoting the storage of past experiences, by facilitating planning based on an evaluation of those experiences, or both. We designed a dynamic spatial task which promotes replay before a memory-based choice and assessed how the content of replay related to past and future behavior. We found that replay content was decoupled from subsequent choice and instead was enriched for representations of previously rewarded locations and places that had not been recently visited, indicating a role in memory storage rather than in directly guiding subsequent behavior.

## INTRODUCTION

Central to cognition, memory allows us to store representations of experience and to later use those representations to inform our future actions. Both of these processes engage a brain structure called the hippocampus (Buckner, 2010; Milner et al., 1998; Squire, 1992) and both have been hypothesized to rely on a pattern of neural activity known as hippocampal replay. During replay, the pattern of neural activity corresponding to a past experience is reactivated in a time-compressed manner (Buzsaki, 2015; Carr et al., 2011; Joo and Frank, 2018). However, the specific role of replay in supporting memory functions, and the principles that determine which experiences are reactivated, remain unknown.

Replay tends to occur during sharp-wave ripples (SWRs), distinctive bouts of high-frequency (150-250 Hz) oscillatory activity in the hippocampal local field potential (Buzsaki, 1986, 2015), and is most often studied in the context of spatial experience. As animals traverse a path through their environment, sets of hippocampal “place” cells are activated in sequence as animals pass through the “place field” of each cell (Foster and Wilson, 2006; Lee and Wilson, 2002; Moser et al., 2015). During sleep and pauses in behavior, time-compressed versions of the same sequences of neural firing are seen, corresponding to a retrieval of activity patterns related to the original experience (Findlay et al., 2020; Joo and Frank, 2018). Replay engages activity across many brain structures, both cortical and subcortical, suggesting that it coordinates a distributed, multimodal representation of experience (Joo and Frank, 2018). During sleep, replay has been linked to memory consolidation, when the repeated activation of hippocampal ensembles is thought to engage plasticity in distributed cortical and subcortical networks to facilitate long-term memory storage (Dupret et al., 2010; Ego-Stengel and Wilson, 2010; Girardeau et al., 2009; Gridchyn et al., 2020; Michon et al., 2019; van de Ven et al., 2016). The role of replay during waking is less clear, however: while awake SWRs have been linked to learning (Fernandez-Ruiz et al., 2019; Jadhav et al., 2012; Nokia et al., 2012), how the content of those SWRs contributes to memory-related information processing remains controversial.

There are currently two leading hypotheses about the role of awake replay. One hypothesis proposes that awake replay serves to retrieve information related to potential future experiences which can then be evaluated to make a choice (Olafsdottir et al., 2018; Yu and Frank, 2015; Zielinski et al., 2020). Under the “planning” hypothesis, the content of replay is expected to predict or otherwise influence subsequent behavior by informing a deliberation process that evaluates replayed options. Evidence for that possibility comes from studies showing enriched replay for the trajectory that a subject is about to take (Pfeiffer and Foster, 2013; Xu et al., 2019) and better coordination of hippocampal-prefrontal activity during replay of the upcoming trajectory in a well-learned task (Shin et al., 2019). Replay of a location associated with an aversive stimulus has also been shown to precede avoidance of that location (Wu et al., 2017), suggestive of a plan for what not to do. Replay has been shown to overrepresent the currently non-preferred or unrewarded option of two possible paths (Carey et al., 2019; Gupta et al., 2010), which could serve to inform the subsequent choice of the alternative, preferred option. Related findings report that replay represents multiple upcoming possibilities (Gupta et al., 2010; Olafsdottir et al., 2017; Shin et al., 2019; Singer et al., 2013), which are also compatible with the idea of replay serving to retrieve possible future options in service of deliberation. Importantly, the planning hypothesis posits that in each of these cases, the specific experiences that are replayed serve as the basis for choice, and thus that replay content influences immediately subsequent behavior, although the nature of that influence (approach or avoid) could depend on what is replayed.

In contrast, many studies report replay content that seems unrelated to upcoming behavior, and thus incompatible with this theory. Replay is most often seen following receipt of reward (Singer and Frank, 2009) and often represents locations leading up to the reward (Ambrose et al., 2016; Diba and Buzsaki, 2007; Foster and Wilson, 2006; Karlsson and Frank, 2009), which can be interpreted as representing the immediate past rather than relating to the future. Replay can also represent experiences that are distant in time and space (Karlsson and Frank, 2009), never-used shortcuts between familiar locations (Gupta et al., 2010), and places that are visible but inaccessible (Olafsdottir et al., 2015). These findings suggest an alternative hypothesis: that rather than informing immediately upcoming behavior, replay instead serves to maintain and strengthen representations of certain places, particularly in the absence of visits to the location (Dudai, 2012; Gupta et al., 2010; Roux et al., 2017; Schapiro et al., 2018). The studies which reported replay biased towards non-preferred task options (Carey et al., 2019; Gupta et al., 2010) are also compatible with the maintenance of nonrecent trajectories. Further corroborating a maintenance rather than a planning role for replay, one of the two studies reported that there was no trial by trial relationship between the replay of the nonpreferred arm and behavioral choice (Carey et al., 2019). In serving a maintenance role, replay would facilitate the long-term storage of particular experiences, but would not necessarily influence immediately subsequent behavior.

Importantly, the behavioral tasks used in previous studies limit the conclusions that can be drawn. Studies that report replay consistent with planning did not include spatially and temporally remote, unavailable, or specific non-preferred options, and thus cannot assess the impact of those options on replay content. Conversely, studies that report replay consistent with memory storage typically use relatively simple tasks with few possible spatial trajectories where planning may be dispensable for accurate performance. Essentially all previously used tasks involve repetitive navigation to a small number of fixed reward locations, making it difficult to disambiguate replay related to prior rewarded experiences from replay of possible future experiences. Moreover, the vast majority of previous studies do not relate replay to behavior on a trial-by-trial basis.

We therefore designed a dynamic memory task for rats that addresses these limitations. This task allowed us to measure whether and how replay content preceding a choice point was related to past experience and to the subsequent choice. Additionally, we incorporated dynamic goal locations in order to assess the influence of reward history on replay content. We then recorded neural activity from the hippocampus during behavior and employed powerful state-space decoding algorithms that allowed us to quantify replay in each subject individually and on a trial-by-trial basis.

Our results revealed a striking lack of correspondence between replay content and subsequent choice. Replay neither increased nor decreased the tendency of the subject to visit the replayed location, regardless of reward history or visit recency of the location. Instead, we found that replay was dominated by two categories of past experiences: places which reliably delivered reward (even after they no longer did so) and places that had not recently been visited, strongly suggesting that hippocampal replay supports memory processes by facilitating memory maintenance and long-term storage rather than guiding upcoming behavior.

## RESULTS

### Rats learn to perform a flexible spatial memory task

Our novel task was designed to fulfill several critical criteria: to provide a defined period where planning would be expected to occur, to require a variety of spatially distinct trajectories, to include dynamic goal locations, and to incorporate many opportunities for replay, particularly before the subject must make a memory-guided choice. The resultant paradigm is an eight-arm spatial version of a win-stay lose-switch task (Figure 1A). The overall task requires the subject to first identify the current goal location (goal arm), continue to choose that arm until the goal changes, and then find the new goal arm. Each self-paced trial in the task begins with the rat triggering the illuminated home port in order to receive a small amount of liquid food reward. After the home port has been triggered, one of the two central ports will illuminate, randomly chosen on every trial. The rat must insert and maintain its nose inside the lit port for a variable duration (2-20 s), after which point a tone sounds and a small amount of reward is delivered. Having the center portion of each trial randomly chosen between two different ports and enforcing an unpredictable delay ensures that animals had to remember a specific goal arm rather than adopt a fixed motor sequence. Following the delay, the outer arm ports then all illuminate, and the rat must proceed to visit one. At any given time, only one outer arm port will provide a large reward (the goal arm), while all other outer arm ports will provide nothing. After visiting a single outer arm, the rat must return to the now-illuminated home port to initiate the next trial. Any port visit that deviates from the defined trial structure causes a “timeout” and requires the subject to restart the trial at the home port.

**Figure 1.**
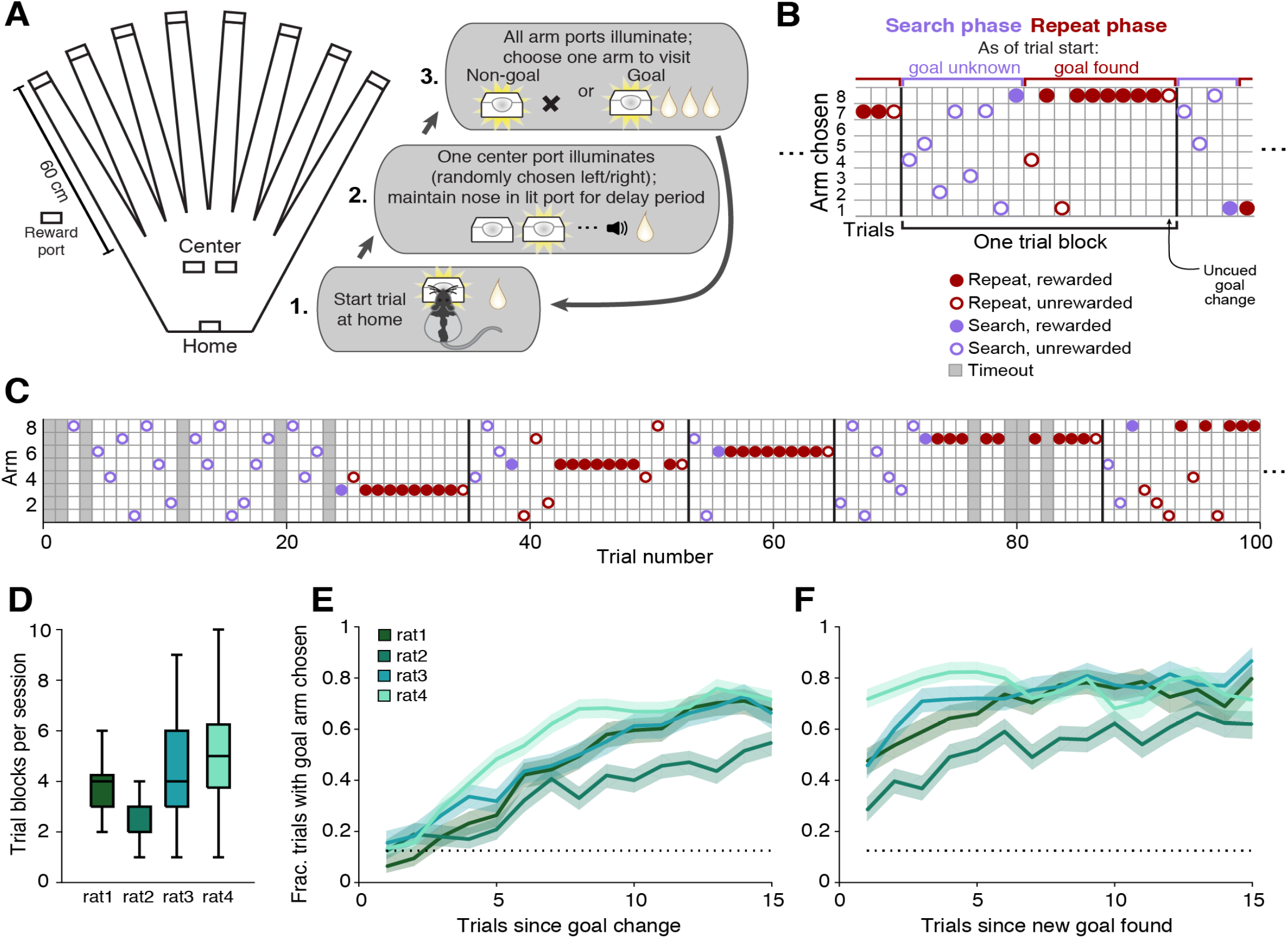
Rats learn to perform a flexible spatial memory task. (A) Overhead view of the track layout, consisting of eight outer arms ending in reward ports and a central “box” area containing the home and center ports. Each trial consists of the three steps illustrated. (B) Schematic illustrates the arm choices of the rat across the two trial types, search and repeat, that make up each block of trials. (C) Expanded series of trials from an example behavioral session including more than 4 complete trial blocks. Trials which contain any deviation from proper visit order (timeouts; grey shading) are excluded from analysis. (D) Boxplot of trial blocks completed per behavioral session for each subject; n = 24, 32, 23, and 33 sessions per subject, respectively. (E) Average outer arm reward rate of the 4 subjects aligned to the first trial after the goal location has changed. (F) Average outer arm reward rate aligned to first trial after first reward is received at a new goal location. For (E) and (F), dotted line indicates chance performance level of 0.125 corresponding to a random choice out of eight possible arms. Shading indicates binomial S.E.M. across trial blocks; n = 89, 85, 105, and 166 trial blocks per subject, respectively.

During the “search phase” (Figure 1B), the rat generally visits different arms on each trial until the goal arm is found. The search phase facilitates frequent exploration of many (eight) arms, providing opportunities to compare replay of experiences across a wide range of visit recencies. Then, during the “repeat phase”, subsequent visits to the goal arm continue to be rewarded until a certain number of rewards have been earned (4-15 rewards; Supplementary Figure 1A and see Methods). The goal location then changes to a new arm that has not previously been the goal arm in the current behavioral session. The subject discovers the uncued change only when it visits the arm that had been the goal and fails to receive the expected reward. The rat then begins a new search phase and the pattern repeats. Together, the search and repeat phases for a given goal location are considered one “trial block”. Over the course of a full behavioral session (60-90 mins), subjects experience several trial blocks (Figures 1C and 1D).

The structure of this task allows us to separate replay preceding the arm choice from replay after the outcome of that choice. The multiple enforced pauses at the home and center ports increase the opportunities for replay before the subject must choose which arm to visit. These periods allow us to determine whether replay preceding a choice is related to the specific choice made. The search and repeat phases were designed to test for different types of planning and potentially different engagement of planning-related neural activity. During the search phase, when the goal is unknown, replay of the previous reward location (previous goal) could help to “remind” the subject that it needs to search elsewhere. Alternatively, planning-related replay might be absent during the search phase due to the lack of a known goal. In contrast, during the repeat phase, replay of the current goal location could inform choices to approach that goal. Across both phases, we can compare the subject’s behavior on trials where we see replay consistent with planning to trials when such replay is absent, allowing us to assess whether the presence or absence of replay of specific locations shapes the upcoming choice. Conversely, pre-choice replay unrelated to subsequent choice but biased toward specific salient locations would provide evidence in support of the memory maintenance and storage hypothesis. Further, replay that is indistinguishable across correct and error trials would also contribute evidence to a maintenance role for replay.

We trained four adult Long-Evans rats to perform the task before implanting a bilateral 30-tetrode microdrive array (Roumis, 2017) targeting the dorsal CA1 region of the hippocampus. We focused on well-trained task performance, rather than initial task acquisition, to have the opportunity to link replay content with reliable behavior and isolate the function of replay during task performance. Thus, although they were continually adapting to dynamic goal locations, our subjects were familiar with the environment and task structure, and had repeatedly received reward from all possible locations. Aligned to the first search trial of each trial block, the reward rate at the outer arms for all subjects began at chance (0.125) and rose (Figure 1E). The speed of this rise reflects the number of trials taken during the search phase of each trial block to find the new goal location and reliably return on subsequent trials. When aligned to the trial after the first reward has been received at a new goal location, the outer reward rate was well above chance, indicating that all subjects preferentially visited the goal location consistently after finding it (Figure 1F). Thus, rats learned to perform this task as intended, alternating between phases of searching for a new rewarded location and repeating visits to a remembered goal. As a key goal of our study is to describe relationships between behavior and replay content that are evident within and consistent across individual subjects, we provide quantification by subject throughout.

### Most SWRs contain interpretable spatial content

As replay events tend to coincide with SWRs, we first identified SWRs using a permissive threshold to ensure that we were examining a large fraction of the actual replay events in the dorsal hippocampus (see Methods). Additionally, we verified that our conclusions were unchanged by using a multiunit-based event detection strategy (Davidson et al., 2009; Pfeiffer and Foster, 2013) instead of a SWR-based one (see Methods). SWRs occurred primarily during reward consumption at each of the stages of each trial (Supplementary Figure 1B). Before the choice point, they occurred during reward consumption at the home port (minimal reward provided) and during the delay period and subsequent reward consumption at the central ports (minimal reward provided). After the trial outcome, they occurred during reward consumption at the current goal arm (large reward provided). As expected, SWRs were rarely detected at non-goal outer arm ports, which did not provide reward (Ambrose et al., 2016).

To assess spatial replay content, we used a clusterless, state space decoding algorithm (Deng et al., 2015; Denovellis et al., 2019; Denovellis et al., 2020). For simplicity and computational efficiency, we first linearized the 2D maze environment into nine 1D segments: the central “box” segment and eight arms (Figure 2A). Our clusterless decoding algorithm incorporates two main components: a marked point process encoding model relating spike amplitudes during movement times (>4 cm/s) to linearized position, and a state space movement model that allows for both smooth, spatially continuous changes in estimated position over time as well as spatially discontinuous jumps between distant locations (Denovellis et al., 2019; Denovellis et al., 2020). The model generates a joint posterior probability of position over the two movement states, thus providing both an estimate of position and an estimate of the movement state for each time bin. The combination of using clusterless data (which incorporates all spikes, not only those from putative single units), the flexible state space movement model, and extremely small time bins (2 ms) allows us to decode changes in the neural representation of position that evolve over a wide range of speeds, including during navigation through the environment and during high speed, temporally compressed replay events.

**Figure 2.**
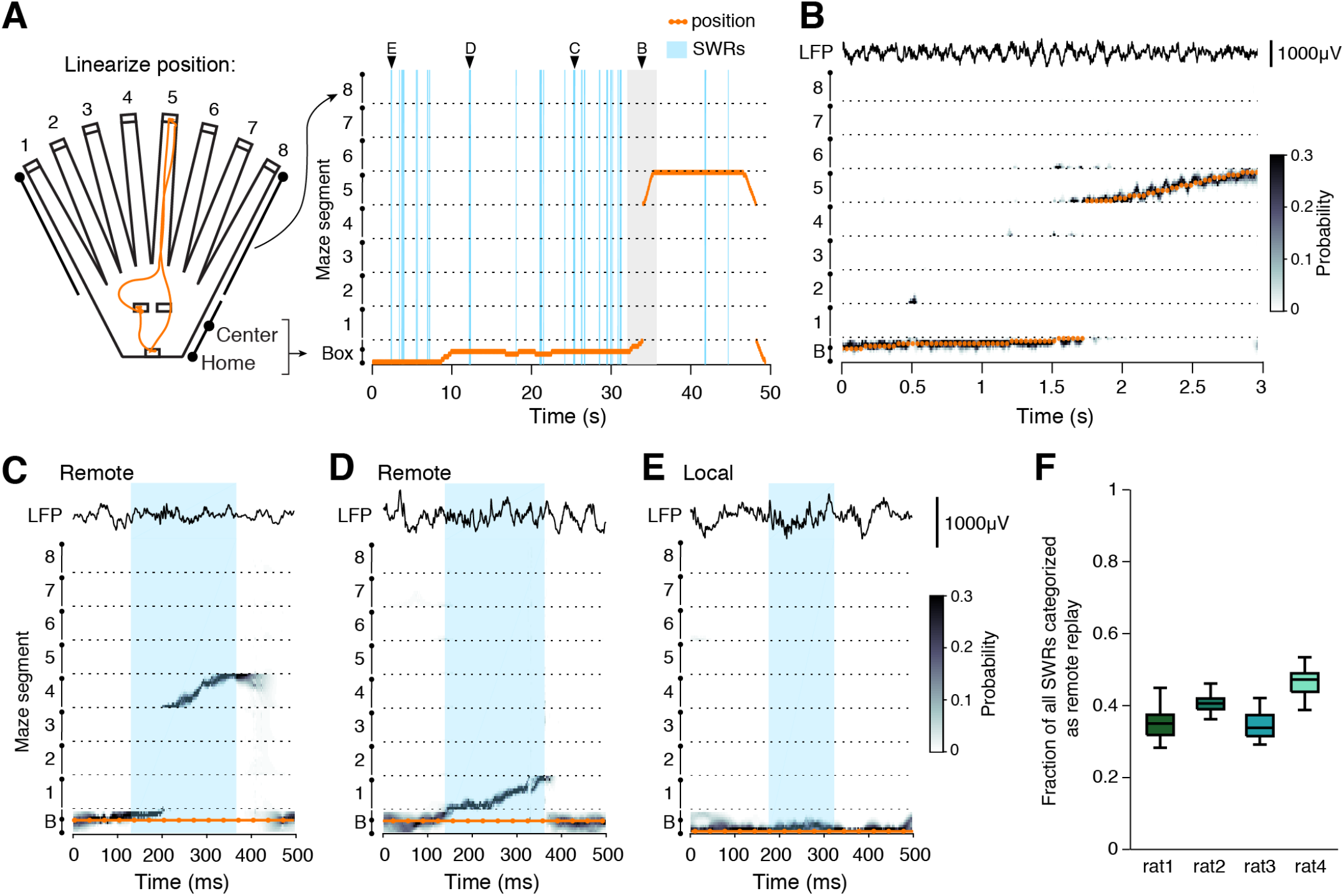
Decoding replay content during SWRs. (A) 2D position is linearized by converting each maze area into a linear segment. Orange trace indicates the 2D (left) and 1D (right) position tracking for a single trial; linearized maze segments are separated by dashed lines. SWR events are indicated in blue. Arrowheads indicate the example SWRs shown in panels (C-D) and the example movement period (grey shading) shown in (B). (B) Decoded position for the movement period as the animal runs from a center port to the end of arm 5. Representative CA1 LFP trace is shown above. (C, D) Two examples of spatially continuous remote replay events occurring during the single trial illustrated in (A); SWR detection boundaries are indicated with blue shading and representative CA1 tetrode LFP is included above. (E) Example of a spatially continuous local event in which decoded position is the same as the actual position. (F) Quantification of the fraction of all detected SWRs that are categorized as remote (spatially continuous with the majority of posterior density in a segment other than the animals’ current location and exceeding 30% posterior density in a single maze segment). Boxplot range indicates variability in remote event fraction across sessions; n = 24, 32, 23, and 33 sessions per subject, respectively.

We first verified that our decoding algorithm accurately tracked the position of the animal during movement (Figure 2B). The median deviation between predicted and real position was less than a single spatial bin (5 cm), indicating that the maximum posterior and real position most often corresponded to the same spatial bin, and this was consistent across all segments of the maze (Supplementary Figures 2A, 2B, and 2C).

Next, we examined the predicted position during times of immobility, specifically during SWRs. We find that the vast majority of SWRs (∼95%) contain spatially continuous representations of the environment (See Methods; Supplementary Figure 2D), including events representing both remote areas of the maze as well as places near the animal’s current position (Davidson et al., 2009; Denovellis et al., 2020; Karlsson and Frank, 2009; Yu et al., 2017). This fraction is much larger than reported in previous studies because our state space algorithm can identify events as spatially continuous that would have been excluded by previous approaches (see Denovellis et al., 2020 for similar results from a different task). We classified spatially continuous events as either remote (majority of probability density in a maze segment other than the one currently occupied by the animal), or local (majority of probability density in the maze segment currently occupied by the animal) replay (Supplementary Figure 2E). Examples of remote and local replay are shown from a single trial of one subject in Figure 2C-E, and for remaining subjects in Supplementary Figure 3. We also verified that our results were similar when using a more permissive definition of remote events (see Methods).

Across all animals, we found that a substantial fraction of detected SWRs contained remote replay: 35.1%, 41.1%, 34.5%, and 47.2% for the 4 subjects (Figure 2F), comparable to prior studies (Davidson et al., 2009; Karlsson and Frank, 2009; Pfeiffer and Foster, 2013). We also note that some previous studies have further characterized the directionality of replay events as forward or reverse. As we were primarily interested in assessing the spatial locations represented across as many events as possible, we did not subdivide events into these categories.

### Replay content is enriched for previous goal locations

These data allowed us to systematically assess the relationship between replay content and behavior. The large majority of spatially continuous replay events occur in the box area, before the subject must choose an arm, and comprise both local and remote representations (Figure 3A). The planning hypothesis predicts that replay before the subject chooses an arm should predict or relate to the subsequent choice of arm; thus, we first focused on understanding the remote events in the box area, which represent specific arms of the maze. To measure the relationship between replay content and both past and future choices, as well as other potentially important locations, we defined four task-relevant arm categories for each trial: the arm the animal chose on the prior trial (past), the arm that the animal will choose on the current trial (future), the current goal arm (current goal; only relevant during repeat phase trials), and the arm that served as the goal arm during the previous trial block (previous goal). Beginning with search phase trials, we asked whether we saw enriched replay of locations based on their category. To ensure that we could assign a single arm to each category, we examined the large subset of search trials in which the past, future, and previous goal arms were all distinct arms of the maze. For completeness, we also quantified the average proportion of events per arm that were representative of the other five arms during these trials (other). We compared the fraction of remote events that corresponded to each category to a null distribution which chose arms randomly from all eight possible options but matched the number of remote events per subject (Figure 3B).

**Figure 3.**
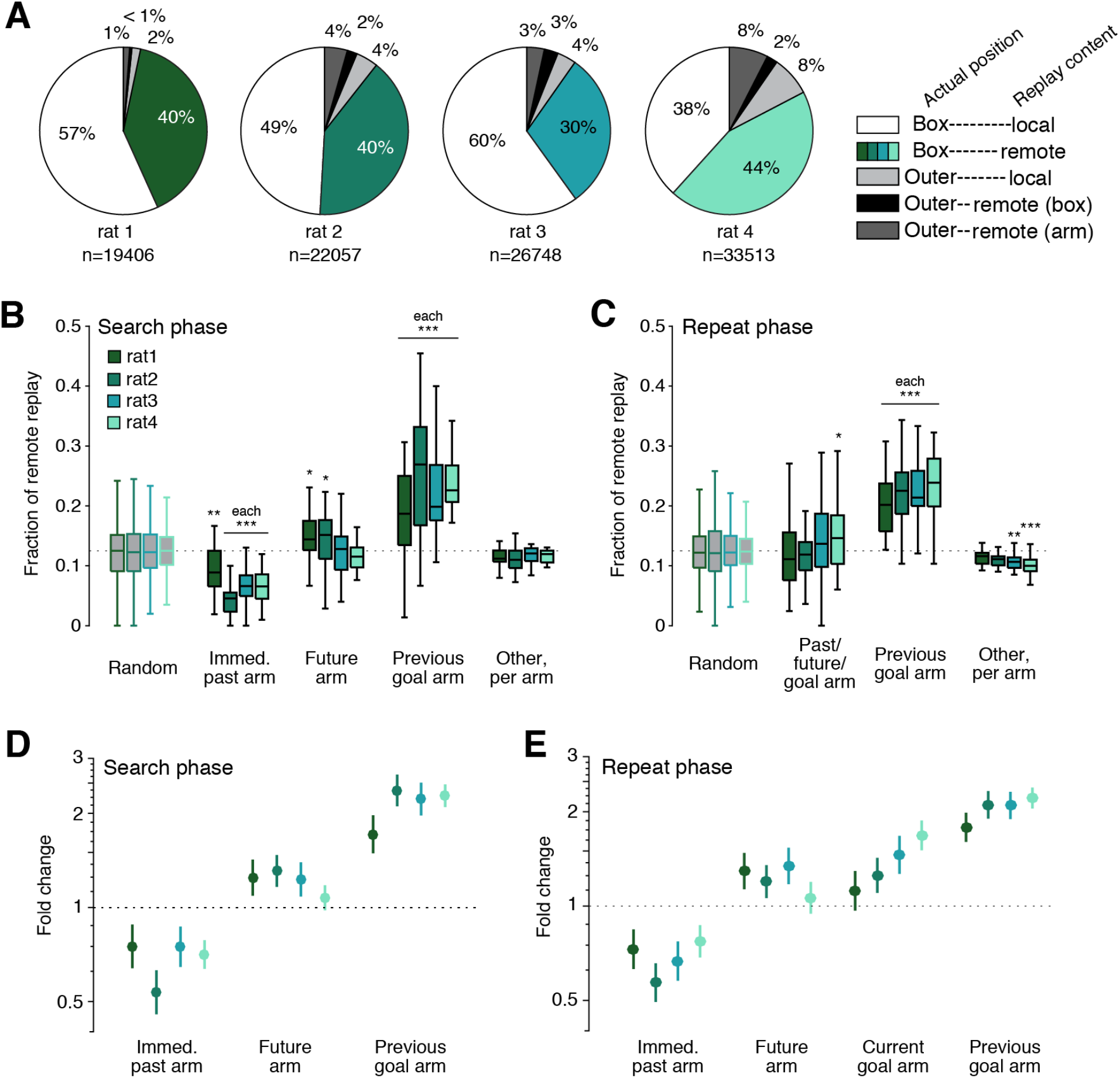
Replay is consistently enriched for previous goal arms. (A) Pie charts for each subject illustrate the proportion of spatially continuous replay events that occur in the box or at arm ports by their content. (B) Boxplots of the fraction of remote replay events in the box area that represent the past, future, previous goal, or other arms during a subset of search phase trials in which past, future, and previous goal arms are distinct. Boxplot range indicates the variability in remote event fractions across sessions; only sessions with at least 5 trials that met the distinct arm criteria were included. N = 22, 23, 16, and 26 sessions per subject, respectively. Significance is calculated relative to a random distribution which is matched for remote event numbers per session for each subject using Wilcoxon rank-sum test. * indicates p<0.05; ** indicates p<0.01; *** indicates p<0.001. (C) As in (B) but using a subset of repeat trials when the past, future, and goal arms are the same and the previous goal arm is distinct. N = 23, 30, 22, and 31 sessions per subject, respectively. (D) Fold change of remote replay rate given arm identity of the past, future, or previous goal based on GLM model. Lines represent 99% confidence interval; n = 711, 848, 725, and 1065 trials per subject; 8 entries per trial. (E) As in (D), but for repeat trials; n = 1347, 1661, 1154, and 1730 trials per subject; 8 entries per trial.

The simplest version of the planning hypothesis suggests that we would see consistent enrichment of replay representing the upcoming choice. That was not the case; while we did see a small but significant enrichment for the upcoming arm in two out of four subjects, this enrichment was not consistent across animals (Figure 3B, future). Our results were also difficult to reconcile with a simple role for replay in consolidating recent past experience: we found far fewer replays of the immediate past arm than would be expected by chance (Figure 3B, past). In contrast, we saw consistent and strong enrichment for replay of the previous goal arm: such replay accounted for roughly 20% of all pre-choice remote replay events during these search trials (Figure 3B, previous goal), even though the previous goal location had not been visited for at least one trial and was not about to be visited (in this trial subset, the previous goal was not the past or future arm). Finally, replay of other arms occurred at roughly chance levels per arm (Figure 3B, other), but considering the number of other arms in this subset of trials (five arms), this indicates that a large fraction of replay events represented arms corresponding to the “other” category.

Perhaps replay during the search phase was enriched for the previous goal rather than the upcoming trajectory because the new goal location was not yet known. To address this possibility, we next looked at pre-choice remote events during the repeat phase, when subjects were reliably visiting the goal arm. However, despite the clear shift in behavioral choices from search to repeat phase, we saw essentially identical results in terms of replay content. During the large subset of repeat trials in which the past, future, and goal arm refer to the same location, distinct from the previous goal arm, all four animals still showed a strong and highly significant tendency to replay the previous goal (Figure 3C, previous goal). In contrast, replay of the future arm still showed only a small and variable effect, despite now also serving as both the past arm and current goal location (Figure 3C, past/future/goal). Thus, even when the goal arm was known, replay of the upcoming arm was inconsistent and was not the main driver of replay content. Instead, the replay of the previous goal location showed the strongest enrichment.

We then extended these analyses to the full set of trials. We assessed the relative impact of each arm category by using Poisson generalized linear models (GLMs) to relate the number of times an arm was replayed to whether that arm was the past, future, current goal, or previous goal arm. We applied these models to the full dataset of either search or repeat trials (see Methods and Supplementary Figure 4). The resultant models include an intercept term that accounts for the overall rate of replay for each subject (not shown) and coefficients that estimate the magnitude of change in replay rate caused by an arm being in a particular category. Because the model posits a linear influence of each predictor on the log of the rate of replay, we converted the resulting coefficients for each predictor into a measure of fold change, which captures the extent to which replay of an arm is amplified or diminished based on its category.

The GLM analyses confirmed and extended our previous findings. During search trials, previous goal arm status had the strongest impact on replay, such that if an arm was the previous goal, there was an approximately two-fold increase in replay rate from baseline (Figure 3D). The past arm showed a significant and consistent reduction in the rate of replay. The future category showed a small increase in replay rate that was significant in three of the four subjects but not in the fourth, corroborating the inconsistent effect seen in the trial subset analyses. Applying the GLM approach to repeat trials yielded similar results (Figure 3E), confirming that for both search and repeat phase, the previous goal arm was the strongest and most consistent driver of replay content, while the replay of the past arm was reliably reduced.

### Replay of current goal location increases with experience

Previous studies reporting replay of upcoming trajectories observed this result in scenarios when the upcoming trajectories led to a predictable, familiar goal location (Pfeiffer and Foster, 2013; Xu et al., 2019). We wondered whether such replay could have been explained by the status of this location as a reliable, repeatedly rewarded goal location rather than as the upcoming choice. If so, then we would expect that in our dynamic task, any consistent replay of the current goal might arise only late in the repeat phase of each trial block, after that arm had been rewarded multiple times.

Our data provided strong support for that explanation. We split repeat phase trials into two groups: those in which the subject had received fewer than five rewards at the current goal location (“early”), and those in which the subject had received five or more rewards at the goal location (“late”). We fit separate GLMs to measure the effect of arm categories on the rate of replay for early (Figure 4A) and late (Figure 4B) repeat phase trials. Indeed, we found a consistent enhancement of current goal replay only in the late portion of the repeat phase. In contrast, the three other categories did not differ substantially between early and late conditions. To measure this effect with finer temporal resolution, we quantified the mean replay rate of the current goal arm for correct repeat trials grouped by the number of rewards received at the goal location (Figure 4C). This analysis revealed a consistent positive relationship between reward experience and replay of the goal location. Quantified on a trial-by-trial basis, we found significant positive correlations between the number of replays of the current goal and the number of rewards already received at that location, although there was substantial variability across trials, as indicated by the low R^2^ values (R^2^=0.033, 0.015, 0.012, and 0.025; p=1.54e-6, 0.0003, 0.0004, and 2.87e-11; n=923, 1007, 847, and 1340 trials per subject, respectively). Together, these findings indicate that replay of the current goal location increases gradually as the subject accumulates reward experience of the location.

**Figure 4.**
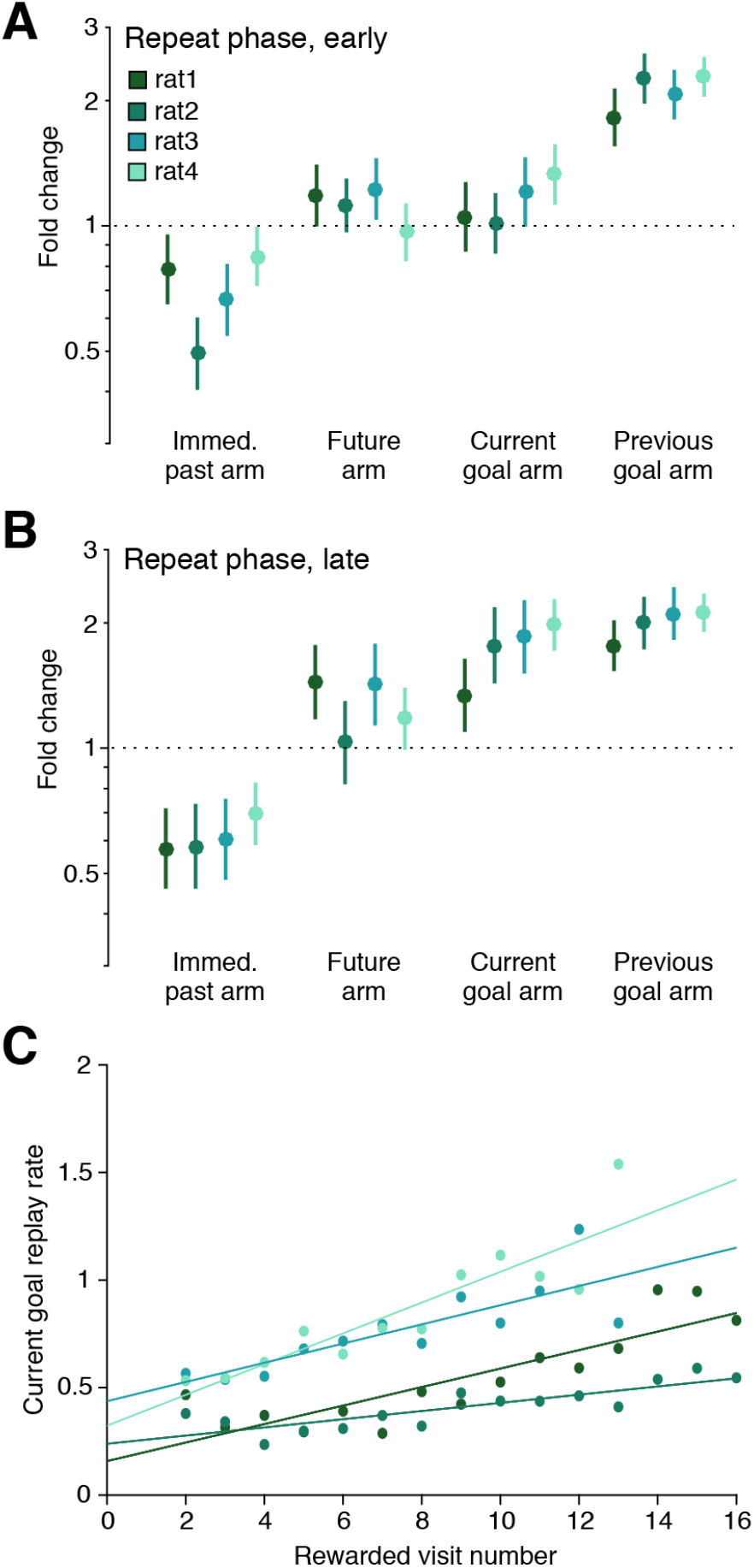
Replay of current goal increases with experience. (A) Fold change of remote replay rate given arm identity of the past, future, current goal or previous goal based on GLM fit to repeat trials that occur before the subject has received five rewards at the current goal location. Lines represent 99% confidence interval; n = 594, 801, 573, 793 trials per subject; 8 entries per trial. (B) Same as (A) for trials when the subject has earned five or more rewards at the current goal location. N = 801, 895, 600, and 967 trials per subject; 8 entries per trial. (C) Replay rate of the current goal on correct repeat trials stratified by the amount of reward experience the subject has had at the current goal location. Points represent an average of all trials with the indicated reward history; only groups containing at least ten trials are plotted. Range of trials per point: 16-90, 33-87, 10-106, and 13-171 per subject, respectively.

### Subsequent arm choice is not biased by replay of future arm, current goal, or previous goal

Our results showed that, on average, the arm that will be visited next is not consistently preferentially replayed during either the search or repeat phase. However, the possibility remained that perhaps replay of the future arm, or the more prominent replay of the previous goal and current goal, could still contribute to shaping upcoming choices. To assess this possibility, we examined the relationship between replay and subsequent arm choice on a trial-by-trial basis.

We first quantified the prevalence of future arm replay. For all subjects, replay of the future arm occurred on average less than once per trial, and was also less prevalent during the repeat phase than during the search phase (Figure 5A). Thus, many trials did not include any replay of the upcoming arm. We next examined error trials during the repeat phase, when the subject has experience with receiving reward at the current goal arm but chooses to visit a different arm. We reasoned that if future replay critically informed behavioral choice, error trials could be related to a lack of sufficient future replay. However, we found no difference in future replay rate between correct and incorrect repeat trials for two out of the four subjects (Figure 5B). For the other two subjects, future arm replay was actually higher, on average, for error as compared to correct trials, the opposite of what the planning hypothesis would predict. This also suggests that it is very unlikely that the error was related to insufficient replay of the upcoming arm.

**Figure 5.**
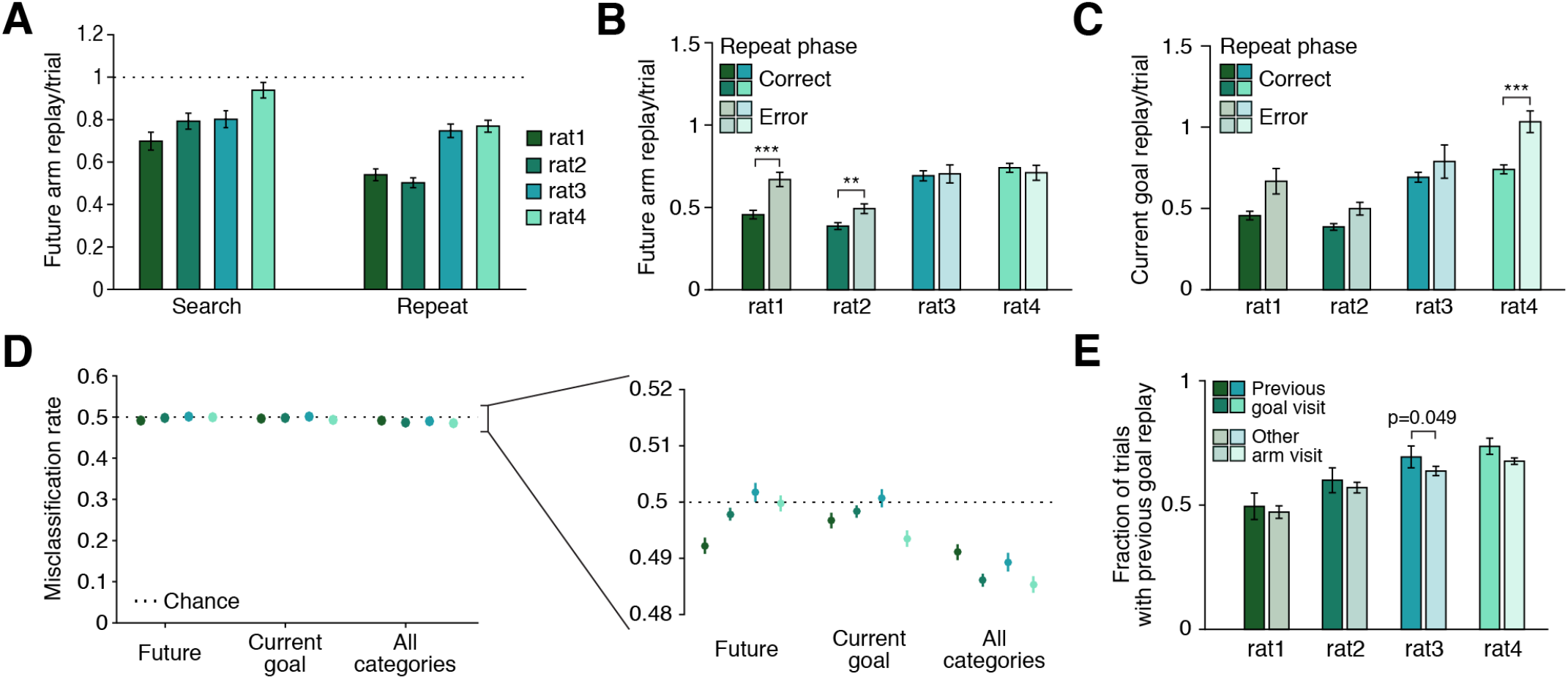
Upcoming behavior is unrelated to replay of the future, current goal, or previous goal. (A) Future replay per trial tends to occur less than once per trial, and more often during search trials than repeat trials for three of the four subjects. N = 534, 617, 570, and 879 search trials per subject, respectively; n = 924, 1019, 894, 1302 repeat trials per subject, respectively. Search vs repeat rate p = 0.001, 5.002e-11, 0.264, and 4.111e-5 for each subject, respectively, using Wilcoxon rank-sum test. (B) Rate of future arm replay during the repeat phase on correct (solid) compared to error (shaded) trials. (C) Rate of current goal replay during the repeat phase on correct (solid) and error (shaded) trials. For (B) and (C), n = 923, 1007, 847, and 1340 correct trials and 424, 654, 307, and 390 error trials per subject, respectively. (D) Cross-validated predictions of correct or error trials are at or very near chance levels. For each subject, a five-fold cross-validated binomial GLM is fit and used to predict correct and error trials based on the number of future replay events, the number of current goal replay events, or the number of events in future, past, current goal, and previous goal categories. Trials are subsampled to match the numbers of correct and error trials such that chance classification rate is 50%; n = 848, 1308, 614, and 780 trials per subset per subject, respectively. Inset is an expanded view around 50%; lines represent 99% confidence intervals reflecting the variability over 1000 subsamples drawn from the full dataset. (E) Fraction of all trials which include replay of the previous goal preceding a visit to the previous goal arm (solid) or preceding a visit to any other arm (shaded). N = 23, 32, 23, and 33 sessions per subject, respectively. Error bars for all panels represent S.E.M.

Replay of the current goal was similarly unrelated to subsequent behavior. We reasoned that current goal replay might be used to guide behavior and thus might differ on error compared to correct repeat trials. Again, we found no consistent significant difference between the amount of current goal replay on correct compared to error trials for three subjects (Figure 5C), and the fourth subject showed a significant increase, rather than decrease, in current goal replay on error trials. This further suggests that the errors could not be attributed to a lack of replay of the current goal arm. Importantly, the overall similarity between replay of the future and current goal on error compared to correct trials, and the subject-specific increase in such replay on error trials was consistent for trials that occurred both early and late in the repeat phase of the trial block, indicating that familiarity with the goal location did not modulate the effect (Supplementary Figure 5A-D).

We also found that the small differences in replay in correct and error trials could not explain the animal’s behavior. We trained a binomial GLM to categorize repeat trials as either correct or error based on the number of future arm replay events, the number of current goal replay events, or the numbers of replay of all four categories: future arm, past arm, current goal, and previous goal. Trials were subsampled so that the number of correct and error trials were matched, and results reflected the mean cross-validated misclassification rate over 1000 subsamples (see Methods). In all cases the models performed at or very close to chance levels, far worse than the animals (Figure 5D). These results, in addition to the small and inconsistent mean differences in replay and the heavily overlapping distributions of numbers of replay events (Supplementary Figure 5A-D) all suggest that differences in replay on error trials are unlikely to directly influence the subsequent error.

We then asked a related question for replay of the previous goal arm: does the robust enrichment for pre-choice replay of the previous goal arm play a role in shaping upcoming behavior by deterring the subject from visiting a no-longer-rewarded location? In this interpretation of planning, replay would serve as an avoidance signal (Wu et al., 2017) and would predict a generally negative relationship between replay of the previous goal and visits to that location.

No such negative relationship was present. Despite its persistent enrichment, we found no evidence that the previous goal replay either deterred or encouraged subsequent visits to the previous goal arm. There was no consistent correlation between the rate of previous goal replay with the number of subsequent visits to the previous goal location during the subsequent trial block (for each subject: R^2^=0.008; p=0.49, R^2^=0.019; p=0.32, R^2^=0.021; p=0.18, R^2^=0.023; p=0.08, respectively). Moreover, if replay of the previous goal was a deterrent, we would expect that on trials with pre-choice previous goal replay, subjects would be less likely to subsequently choose to visit the previous goal location. However, we found that previous goal replay was no more or less likely to precede visits to the previous goal as it was to precede visits to any other arm (Figure 5E). Altogether, our findings are not consistent with replay of the future arm, current goal, or previous goal influencing the upcoming behavioral choice, but instead indicate a decoupling between replay and upcoming behavior.

### Previous goal replay persists through multiple trial blocks

All of our findings above are consistent with an enhancement of awake replay for locations that have been consistently associated with reward. This begins in the late part of the repeat trials and continues when the current goal becomes the previous goal during the subsequent search and repeat phases. We next examined how long such enrichment persists. We calculated replay events per trial for each goal arm as that arm transitioned from rewarded to unrewarded, then computed the mean across all trial block transitions with all replay rate curves centered on the last rewarded trial of the block. (Figure 6A, solid lines). For comparison, for each trial block, we also calculated the mean replay rate across the same window of trials for all arms that were never the goal location during that behavioral session (Figure 6A, dashed lines). In the same manner, we quantified visits to each goal arm over the same trial window, as it transitioned from being rewarded to unrewarded, and compared them to visits to unrewarded arms during this window (Figure 6B).

**Figure 6.**
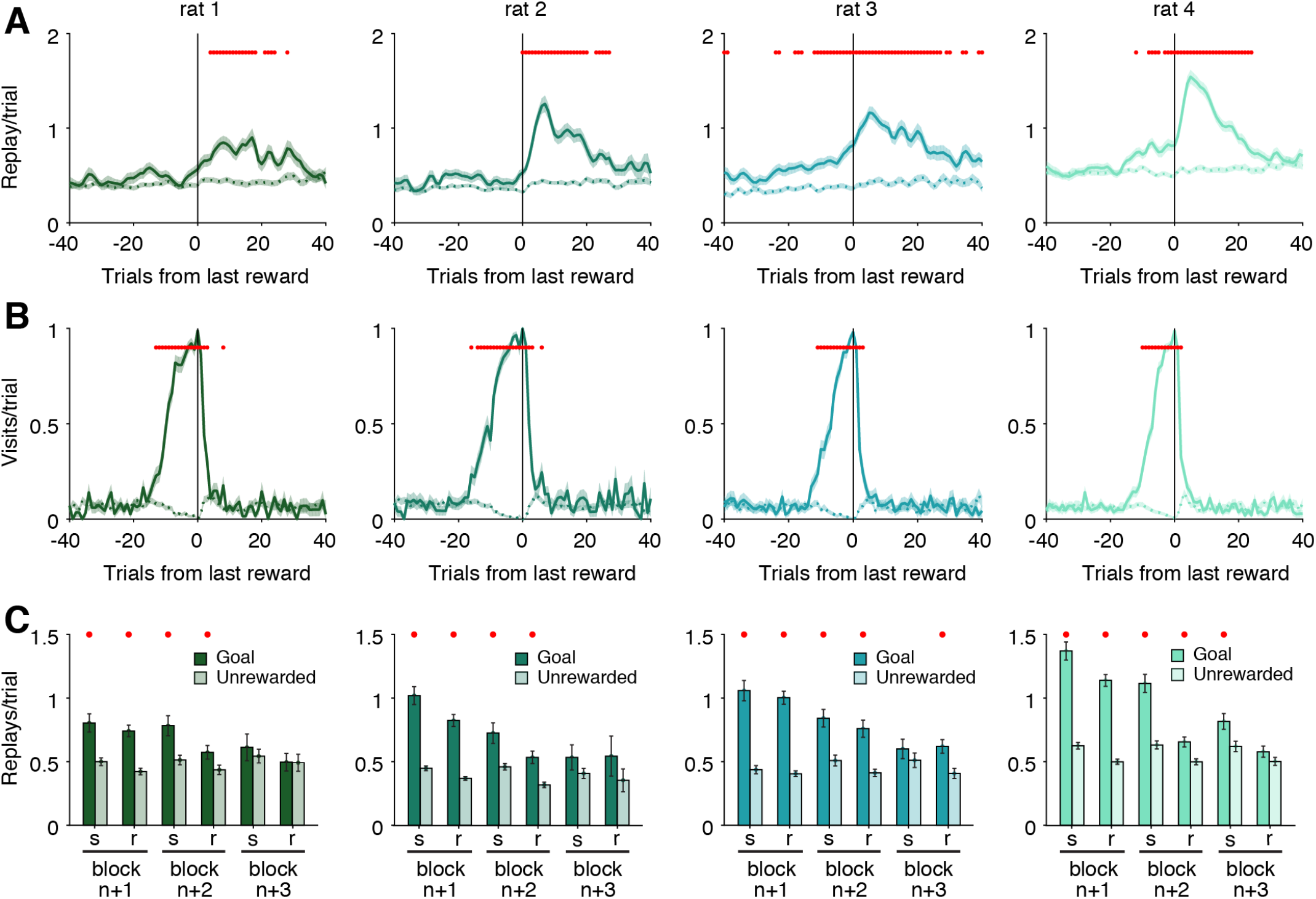
Replay of previous goal continues long after behavior changes. (A) Mean replay rate curve of a particular goal location for each subject aligned to the last rewarded trial of that goal’s block of trials. Solid line indicates replay rate of the goal arm; dashed line represents replay rate of non-rewarded arms during the same span of trials. (B) Mean visit rate to the goal (solid) or non-goal (dashed) arms aligned to the same span of trials as (A). For (A) and (B), shading indicates S.E.M. (C) Mean replay rate of goal and non-goal arms over search (s) and repeat (r) sub-blocks for the three trial blocks after a particular goal location has changed. Error bars represent S.E.M. For all panels, red dots indicate which comparisons exceed a 0.05 significance threshold after Bonferroni correction for multiple comparisons (80 comparisons each for (A) and (B); 6 comparisons for (C)). Data included from 89, 85, 104, 161 goal arms and 347, 437, 268, 397 unrewarded arms per subject, respectively.

These analyses confirmed the dissociation between previous arm replay and previous arm visits. The enrichment for replay of a goal location peaked several trials after it ceased to provide reward and continued for many subsequent trials. This enrichment was extended and significant for all subjects (Figure 6A). We also saw elevated replay before the transition for two out of the four subjects (consistent with replay of the current goal in these subjects shown in Figure 3C). In contrast, behavioral bias to the previously rewarded arm decays extremely quickly, returning to baseline levels within a few trials after the switch (Figure 6B).

Moreover, we also find that previous arm replay continues throughout the second subsequent trial block, when there is yet another new goal arm and the behavioral bias to visit the previous arm is undetectable. We calculated the replay rate of each previous goal arm for the three subsequent trial blocks after it served as the goal, separating search and replay phase for each block. This analysis revealed that the enhanced replay of the previous goal location persisted for two complete trial blocks for all four animals (Figure 6C).

### Pre-choice replay is biased toward non-recent past

The enrichment for representations of the previous goal compared to the reduction of representations of the immediate past arm indicates that not all past experiences are equally prioritized for replay. The search phase of this task provides an opportunity to assess the relationship between recency and replay, while controlling for reward history. Here we leverage the many “other” replay events during search trials – specifically those representing arms that never served as the goal location in the current behavioral session. We found consistently higher replay rates for arms that had not been visited within 5 trials compared to those visited within one or two trials (Figure 7). Although highly variable, the replay rate of arms visited within 1-5 trials was strongly correlated with visit recency (for each subject: R^2^=0.011; p=1.04e-4, R^2^=0.028; p=4.04e-13, R^2^=0.012; p=7.94e-5, R^2^=0.018; p=2.76e-8, respectively), indicative of a consistent bias against replay of more recently visited unrewarded arms. We conclude that pre-choice remote events prioritize non-recent past experiences in addition to previous reward locations.

**Figure 7.**
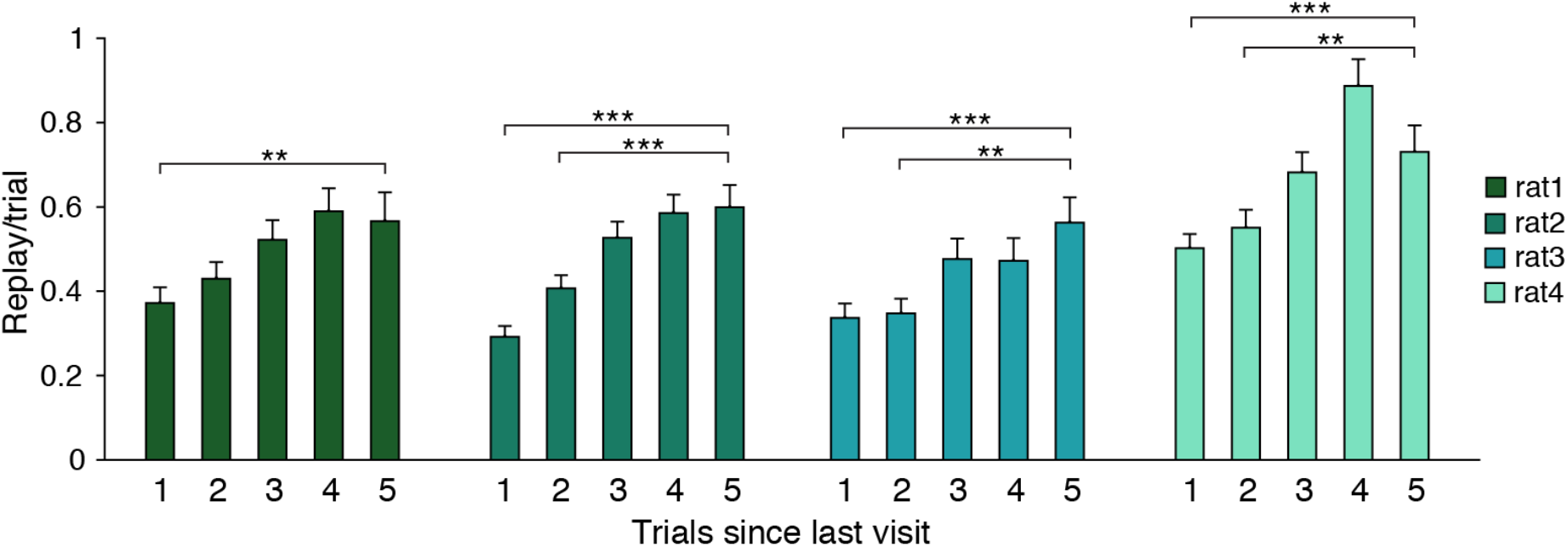
Replay of unrewarded arms is biased toward non-recent past. During search trials, the replay rate of arms that have never been rewarded are categorized based on whether they were visited 1, 2, 3, 4, or 5 trials ago. Error bars indicate S.E.M. over trials; statistical comparisons were calculated between the five trials ago condition and each other bar. * indicates p<0.05; ** indicates p<0.01; *** indicates p<0.001; trials per bar range from 159-433, 212-555, 144-386, and 198-520 trials per subject, respectively.

### Enhanced replay of previous goal at outer arms

If, as our results suggest, the most parsimonious explanation for the replay content that we observe relates to maintaining representations of important and non-recent experiences, then we might expect to see this not only for pre-choice replays, but also for the smaller number of replays that occur after the choice, in the outer arms. Importantly, these events almost exclusively occur during reward consumption at the current goal arm port because unrewarded arm visits are generally extremely brief and rarely include SWRs. We first quantified the fraction of replay events corresponding to the current arm (the goal arm) or the previous goal arm on rewarded trials where the past arm was the same as the current arm. Given that replay is strongly biased to initiate with representations at the current position of the animal (Ambrose et al., 2016; Diba and Buzsaki, 2007; Foster and Wilson, 2006; Singer and Frank, 2009), we omit the comparisons to a random distribution across the eight arms.

As expected, we saw strong enrichment for the current arm. However, we also observed greater replay of the previous goal compared to other arms (Figure 8A). We used a Poisson GLM to quantify these effects during all rewarded trials and found that in addition to overrepresentation of the current arm (the goal arm), we also saw consistent, strong enrichment for replay of the previous goal arm (Figure 8B). These results demonstrate that the enrichment for replay of the previous goal arm is not just specific to pre-choice replay but is evident throughout the task.

**Figure 8.**
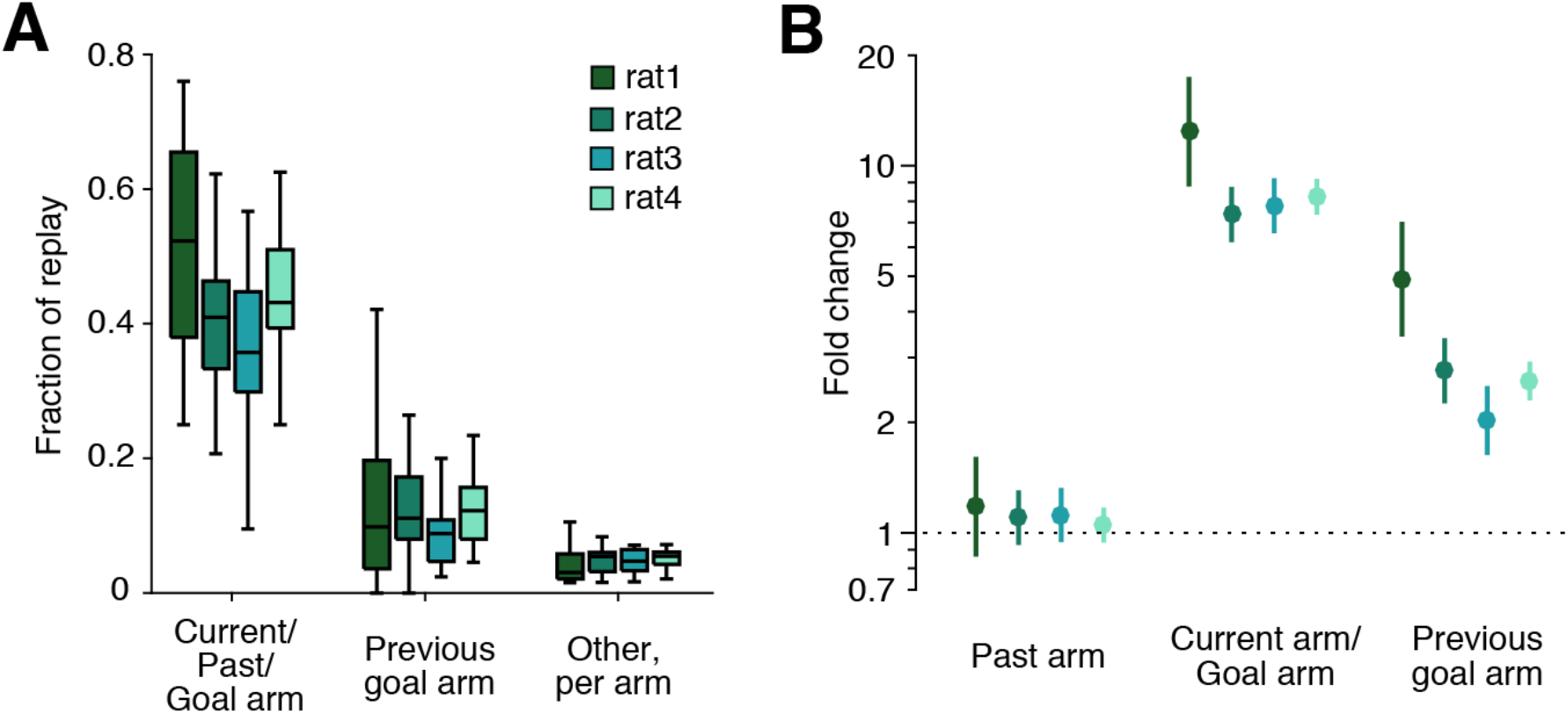
Replay at the outer arm ports. (A) Fraction of non-box events at rewarded outer arm ports that correspond to the current arm/past arm/goal, the previous goal, or any other arms during the subset of trials when the past and current arm are the same and distinct from the previous goal arm. N=333, 958, 1161, 3316 events per subject, respectively. (B) Fold change of arm replay rate given arm identity of current arm/past arm/goal, the previous goal, or any other arms based on GLM model. Lines represent 99% confidence interval; n=948, 1026, 868, 1372 trials per subject; 8 entries per trial.

## DISCUSSION

We investigated the role of awake replay in guiding upcoming behavior in the context of a spatial memory task designed to provide ample opportunity for such replay. This task provided several key advantages over previous paradigms. The task structure separated replay preceding a choice from replay after the outcome of that choice, providing distinct periods in which to look for planning-related replay. Further, the incorporation of changing goal locations and regular searching across many arms allowed us to disambiguate the distinct effects of reward history and visit recency on replay. Our results link replay not to immediate future choices but instead to reliably rewarded and less recently visited locations, consistent with a role in memory maintenance and storage.

The hypothesis that pre-choice replay serves to guide upcoming behavior predicts that the content of replay would be related in some way to the choice that followed. In the simplest case, replay content might be dominated by representations of the upcoming choice, such that the strongest signal conveyed to downstream brain areas would represent the plan for upcoming trajectory. Our results are incompatible with such a model: replay was not consistently enriched for representations of the upcoming arm. Further, we did not find any difference in future arm or current goal replay between correct and error trials, indicating that errors were unlikely to result from any alteration in replay content.

A more complex but more plausible version of the planning hypothesis proposed that replay events provide internal mental exploration of multiple future options (Olafsdottir et al., 2018; Yu and Frank, 2015; Zielinski et al., 2020). Under this hypothesis, some other brain region evaluates each option and then makes a decision, leading to the behavioral outcome. Such deliberation would be expected to engage representations of multiple possible options, likely including the ultimately chosen trajectory. Trial-by-trial analyses demonstrated that many trials did not include any replay of the future arm, and that future arm replay occurred less frequently per trial than replay of the previous goal or other unrewarded arms, indicating that it would be an unreliable and noisy guide for upcoming behavior. Further, error trials were not consistently different from correct trials in terms of future or current goal replay rate, and in fact trended toward having more, rather than less, of both types of replay. Even replay of the previous goal, the most enriched category, did not affect the frequency of subsequently visiting or avoiding the previous goal arm. Overall, we find no evidence that replay of a particular arm increases the frequency that the animal will either visit or avoid that arm. Our findings are thus difficult to reconcile with a model that posits that replay events provide the set of options for upcoming behaviors. Indeed, to comply with such a deliberation scenario, the replay we observe would have to represent a combined signal consisting of both planning-related information alongside many other events – the correct parsing of which would require selective routing or differential evaluation in downstream structures. We are not aware of evidence for such content-specific processing of replay events in the brain.

Instead, our findings are consistent with a singular role for replay: to facilitate longer term memory storage biased toward specific experiences. We find that the experiences represented during replay are often places where the subject has received multiple prior rewards, including current goal locations toward the end of trial blocks as well as locations that have recently been the goal, but are no longer providing reward. Additionally, we find that replay often represents locations which have not been visited recently; these representations may be most at risk of degradation, forgetting, or memory interference. These findings are consistent with there being a baseline level of remote replay for all locations in an environment, with an experience-dependent modulation of that baseline that suppresses replay of immediate past experiences and enhances replay of reliably rewarded or otherwise salient experiences.

Overall, our findings argue for a role for replay in maintaining a high-resolution, accurate representation of locations and experiences to reduce the chance of forgetting or memory interference among these representations (Dudai, 2012; Roux et al., 2017; Schapiro et al., 2018). Here it is important to specify what we mean by “maintenance.” For instance, the reinforcement learning field has suggested that replay might serve to update value representations of a particular place. In this case, replay has been predicted to prioritize content that would update value associations to promote optimal behavior, such as places that are likely to be visited or those that have changed value (Mattar and Daw, 2018). However, because the rapid shifts in behavioral strategies between search and repeat phases far outpace the much slower changes in replay content that we see over trial blocks, and because replay content does not relate to behavior on a trial-by-trial basis, replay seems unlikely to implement the value updates that could drive behavior. Instead, the replay of previously rewarded and temporally remote locations likely serves to strengthen the representation of the location – the multimodal combination of sensory features and past experiences that characterize each location in the environment – by facilitating plasticity in distributed cortical and subcortical networks (Dudai, 2012). While this representation may well include some type of reward associations – replay of reward locations has been seen to engage neurons in brain regions associated with rewards and reward expectations (Gomperts et al., 2015; Lansink et al., 2009; Sosa et al., 2020) – it is possible that replay may reinforce current location-value associations without changing the values themselves. Alternatively, if replay does update values, it could be the case that those value associations comprise only a part of the broader set of computations that drive behavior in our task. Consistent with this possibility, one study reports that the extent to which reward-related neurons engaged with replay of reward locations was not predictive of behavioral performance on a trial-by-trial basis (Gomperts et al., 2015).

Nonetheless, for these stored representations to be useful, they must have the capacity to influence some future behavior. Our results argue against a direct, trial-by-trial influence of replay on behavior; instead, this influence may be indirect. We propose that decision-making processes are not directly engaged during replay. Instead, during replay, representations of experiences are strengthened and updated. Later, outside of replay, neural ensembles associated with those representations could be engaged by a planning process that may directly drive subsequent behavior – such as during “theta sequences” where representations of future possibilities can be expressed (Kay et al., 2020; Papale et al., 2016; Redish, 2016; Tang et al., 2021).

Importantly, the idea that replay serves to strengthen representations of specific experiences provides a unifying explanatory framework for the large majority of previous results, including both those linking SWRs to behavior during learning and those describing the relationship between replay and behavior in the context of well-learned tasks. Previous studies which demonstrate behavioral consequences of SWR manipulation (Fernandez-Ruiz et al., 2019; Jadhav et al., 2012; Nokia et al., 2012) and which relate SWR activity to upcoming behavior (Singer et al., 2013) specifically report these effects during task acquisition. During early learning, replay could help “connect the dots” by storing information about the structure of the environment and associating experiences with outcomes (Barron et al., 2020; Wu and Foster, 2014). As these increasingly accurate and refined task representations evolve, they could inform downstream planning processes, and thereby provide an opportunity for replay to more closely relate to immediately subsequent behavior while still serving a memory storage role.

Critically, we can also account for the profound diversity of results from previous reports relating replay to behavior in well-learned tasks. The strongest evidence for replay of the upcoming trajectory come from tasks where that upcoming trajectory leads to a predictable reward location (Dupret et al., 2010; Pfeiffer and Foster, 2013; Xu et al., 2019). Our results suggest that this occurs not because the replay is directly guiding the subject to the reward location, but instead because the subject has received reward many times at this location in the past. Importantly, Xu et al. do report that error trials fail to show the enrichment for replay of the upcoming arm that they see during correct trials in their reference memory paradigm. However, the error trials in this task appear to occur mainly during initial learning of the day’s reward locations; we suggest that error trials show a different pattern of replay because they occur before the goal locations have become predictable, compared to the many successful trials later in the session.

Replay reflecting both upcoming options as well as past options in spatial alternation tasks (Olafsdottir et al., 2017; Shin et al., 2019; Singer et al., 2013) may also be more parsimoniously explained as replay of historically rewarded locations rather than a complex combination of future deliberation and storage of recent experience. We note, however, that Shin, Tang, et. al. (2019) reported that while hippocampal replay content before a memory-based choice was not predictive of upcoming trajectory, hippocampal-prefrontal cortical (PFC) coordination was specifically enhanced during replay of the upcoming, as compared to the alternative, trajectory. This increased coordination was only seen when subjects had fully learned to perform the alternation task, a time when replay is less frequent (in this case, less than once per trial) and may be less critical for behavior (Jadhav et al., 2012; Karlsson and Frank, 2009; Shin et al., 2019; Singer et al., 2013). This leads us to speculate that the coordinated PFC activity during replay may reflect task performance, but not directly drive behavior (Tang et al., 2021).

Results demonstrating replay of non-preferred options (Carey et al., 2019; Gupta et al., 2010) also fit well within our framework, as we would expect to see enhanced replay of the nonpreferred location both because it was a reliably rewarded location and because it has not been visited lately. Finally, observations of replay representing inaccessible locations or trajectories would be predicted by our framework (Olafsdottir et al., 2015); “never visited” is the extreme of “not recently visited” – replay in these scenarios serves as the only way to form and consolidate representations of known but inaccessible paths, as physical experience of these locations is disallowed.

In addition to replay of particular past experiences (previously rewarded locations and arms that have not been recently visited), we also observe frequent replay of the local maze segment, both in the box area of the maze and at outer arm ports on rewarded trials. This large fraction of local events is likely influenced by the initiation bias seen in replay (Davidson et al., 2009; Karlsson and Frank, 2009): approximately 80% of events begin with representations of locations close to the current position and proceed away. Because our maze segments are fairly large, such “local” replay can include both extended representation of single location as well as trajectories that traverse sub-segment distances, such as from the home port to a center port, or along the length of the rewarded arm. Because the subject is receiving reward at all of these locations – small amounts in the box area and a large amount at the goal arm – such replay is consistent with many reports of replay representing the immediate path to reward seen during reward consumption (Ambrose et al., 2016; Bhattarai et al., 2020; Diba and Buzsaki, 2007). This result suggests a distinction between recent rewarded experiences of the current, local, maze segment which are likely to be replayed, and recent unrewarded representations of remote locations, which are unlikely to be replayed.

Finally, we note that this unifying framework provides clear parallels between waking and sleep replay. Studies of SWRs and replay during sleep have long hypothesized that replay serves to consolidate memories by promoting their storage in distributed cortical-hippocampal networks. Our findings are consistent with the hypothesis that replay serves this function during both waking and sleep (Carr et al., 2011; Roux et al., 2017; Schapiro et al., 2018). Awake replay tends to have greater fidelity to the original experience compared to sleep replay (Karlsson and Frank, 2009; Tang et al., 2017) and greater engagement with subcortical dopaminergic populations (Gomperts et al., 2015), indicating that there may be differences in what is being stored (Roumis and Frank, 2015) during the two states. Nonetheless, replay in both states prioritizes particular experiences over others (Carey et al., 2019; Dupret et al., 2010; Girardeau et al., 2017; Gupta et al., 2010; Michon et al., 2019), suggesting that replay is a general mechanism for preserving neural representations of behaviorally relevant experiences.

## Acknowledgements

The authors are grateful to P. Dylan Rich and Tom Davidson for helpful discussion, Brett Mensh for thoughtful editorial suggestions, Alison Comrie, Abhilasha Joshi, and Kyu Hyun Lee for feedback on the manuscript, and all members of the Frank laboratory for support throughout the project. We also thank Alan Kaplan and the Lawrence Livermore National Laboratory High Performance Computing Center for providing computational resources. This work was supported by the Simons Collaboration on the Global Brain (grant 521921 to L.M.F. and postdoctoral fellowship 500639 to A.K.G.), NIH Brain Initiative (K99AG068425 to A.K.G.) and Howard Hughes Medical Institute (L.M.F.).

## Author Contributions

A.K.G. and L.M.F. conceived the study. A.K.G. and D.A.A.M. collected data. E.L.D., D.F.L., M.E.C., D.K.R., and U.T.E contributed to the development of analytical techniques. D.B.K. contributed to the development of the behavioral task. A.K.G. analyzed the data. A.K.G. and L.M.F. wrote the manuscript with input from all authors.

## METHODS

### Animals

Data from four male Long-Evans rats (Charles River RRID:RGD_2308852; aged 4-9 months; 450-550 g) were included in this study. Animals were kept on a 12-hour light-dark cycle (lights on 6am - 6pm) and initially had *ad libitum* access to food (standard rat chow) in a temperature- and humidity-controlled facility. Beginning one week prior to and extending throughout behavioral training and data acquisition, rats were singly housed and food restricted to a maximum of 85% of free-feeding weight. All procedures were approved by the Institutional Animal Care and Use Committee at the University of California, San Francisco.

### Behavioral Training

The behavioral setup included a small sleep box (12” x12” x16” walls) and a maze environment equipped with automated ports. Task control was managed by an Environmental Control Unit (SpikeGadgets) running custom behavioral code written in Python and Statescript (SpikeGadgets). Each translucent custom port (Roumis, 2017) includes an internal white LED, an infrared photogate and a reward delivery tube connected to a syringe pump (Braintree Scientific) which can dispense precise quantities of reward (evaporated milk + 5% sucrose) when an IR beam break is detected. The maze was roughly 1 m x 1 m, surrounded by 16” external walls with transparent internal walls of the same height between arms. From the maze, subjects could see room cues including wall markings, the door, the sleep box, and a desk. The maze environment contained 11 ports (home port, 2 center ports, and 8 arm ports). The home and center ports dispensed 50 μL of milk while outer ports dispensed 150 μL. Subjects underwent approximately three weeks of 2-3 60-min training sessions per day to reach complete task performance. Ultimately, subjects performed the complete task as follows: each self-paced trial was initiated by a nose trigger at the home port and receipt of reward. Subsequently, one of the two central ports would illuminate (chosen randomly), and the rat had to place its nose in the illuminated port and maintain this position for a delay period, after which time a sound cue occurred and reward was delivered from the port. The delay was randomly drawn on each trial from a limited selection spanning ∼2-20 s. After reward consumption at the central port, all arm ports were illuminated, and the rat had to visit a single arm before returning home to initiate the next trial. At any given time, only one arm port (“goal arm”) would provide reward; the others would provide nothing. Any deviation from this pattern of visits resulted in a “timeout” with no port illumination or reward delivery for 25-30 s, after which time the rat had to restart at the home port.

Rats had to sample various arms on early trials until the rewarded location was found, then continue to visit this location on subsequent trials. After a certain number of rewards were earned at the goal arm, a new goal location was randomly chosen from the arms that had not yet served as goal in the current session. Subjects rarely completed trial blocks at all eight arms, but when this occurred, the goal list reset and all arms became valid goal options. All subjects began with 15 rewards per goal location before switching during early behavior sessions, and this was reduced to 10 rewards per goal for later sessions. For two subjects, it was further reduced to a range of 4-12 rewards, with one value from that range selected randomly per behavioral session (Supplementary Figure 1A).

To achieve reliable performance on this task, behavior was gradually shaped using a series of simplified variants. Subjects first completed 3-4 60-minute sessions of training on a walled linear track with a reward port at each end. Motivated subjects (>50 rewards on final linear track session) advanced to learn the spatial memory task. Subjects next acclimated to the maze environment and learned to visit ports in the correct order (home port - lit center port - any outer port) with all outer ports dispensing reward. A “timeout” was triggered by any incorrectly ordered visits; this timeout increased in length from 3 s to 25-30 s over 3-5 days. A delay was gradually introduced at the center ports, beginning with ∼0.5-2 s and gradually reaching ∼3-10 s (3-5 days). Finally, the number of rewarded outer ports was gradually reduced from all eight to a single goal arm (5-7 days). The inclusion of two center ports ensured that the subject could not simply develop and rely on a fixed motor routine to navigate to the goal arm repeatedly. The two ports were approximately 7.5 cm apart, and a small plastic wall prevented the subject from walking over the port; it instead had to choose to navigate around the ports to reach the arms. From the left port, subjects tended to turn to the left and then proceed out to the arms; from the right port, subjects tended to turn right and then walk to the arms. As a result, the heading directions and routes taken to the arms typically differed between left and right center port trials (data not shown).

### Neural Implant

Best-performing subjects were selected for surgery and allowed *ad libitum* food access without behavioral training for at least five days prior to implant. Each implant housed 30 independently moveable nichrome tetrodes (12.5 μm, California Fine Wire), cut at a sharp angle and gold plated to a final impedance of ∼200-350 kOhms. The implant was stereotactically implanted such that each of the bilateral cannulae containing 15 tetrodes (1.5 mm diameter) was centered at 4 mm AP, ±2.6 mm ML relative to skull bregma. A screw placed over cerebellum served as global reference. Subjects were allowed at least five and up to ten days for recovery after surgery, with *ad libitum* food and daily tetrode adjustment until dorsal CA1 cell layer was reached.

### Data collection and processing

After recovery, subjects were reintroduced to the task apparatus and given 3-6 days to regain reliable performance before neural data acquisition began. Each day, subjects completed 2-3 45-90-min behavioral sessions flanked by 30-min sleep sessions in the rest box. Continuous 30 kHz neural data, environmental events (port beam breaks, lights, reward delivery), online position tracking of a head-mounted LED array, and 30 Hz video were collected using a 128 channel headstage and MCU/ECU system (SpikeGadgets). Hippocampal local field potential (LFP) traces were generated for each tetrode by filtering continuous signal from one channel of the tetrode between 0.1-300 Hz and then referenced by subtracting the signal from a tetrode located in ipsilateral corpus callosum. In parallel, the continuous signal was referenced and filtered between 600-6000 Hz, and spike events were detected when the voltage exceeded 100 μV on any channel of a tetrode. Any small (<1 s) gaps in online position tracking data due to camera occlusion or periods of unrealistically high movement velocities were interpolated and the full position trace was smoothed using a Loess method; this smoothed trace was used to calculate velocity.

### Histology

At the conclusion of data collection, tetrode locations were marked with electrolytic lesions and the subject was transcardially perfused with paraformaldehyde 24 h later. The brain was fixed, sectioned into 50 μm slices, and Nissl stained to enable the localization of tetrode tips, as described in (Sosa et al., 2020).

### Analyses

All analyses were performed using custom code written in Matlab 2015b (Mathworks) and Python3.6.

### Behavioral analysis

Trial landmarks were extracted from behavior log files (Statescript; SpikeGadgets) and included trial start and end, the times of each port trigger and any resultant reward delivery, and whether the trial included a timeout (if so, it was excluded from further analysis). Trials were classified as search if the goal arm had not yet been found and included the trial upon which the goal arm was first visited. Repeat trials included every subsequent trial after the first visit to goal, including those that did not include visits to the goal arm (classified as repeat errors), and included the final unrewarded trial of each trial block. For each trial, the arm categorized as “future” was based on the arm port triggered on that trial; any partial detours into other arms that did not include a port trigger were not reflected. The “past” designation was given to the arm port visited on the previous trial, even if this visit occurred during a timeout.

### SWR detection

SWRs were detected using a consensus method based on the envelope of the ripple filtered (150-250 Hz) trace, smoothed and combined across all CA1 cell layer tetrodes (Kay et al., 2016). Events were detected as deviations in the consensus trace exceeding 2 SD above total session baseline (mean) for at least 15 ms. Each event start was defined as the time at which the consensus trace first crossed session baseline before the threshold crossing; event end was the time at which the trace returned to session baseline after the event. SWRs were only detected during immobility times (velocity <4 cm/s). For multiunit activity (MUA) event detection, a histogram of spike counts was constructed using 1 ms bins; all spikes >100 μV on tetrodes in CA1 cell layer were included. The MUA trace was smoothed with a Gaussian kernel (15 ms SD), and the mean and standard deviation of this trace during immobility periods (< 4 cm/s) was calculated (Davidson et al., 2009). Deviations in the multiunit trace exceeding 3 SD above the mean during immobility periods were considered MUA events; event start and end were defined as the times before and after the event at which the trace returned to the mean immobility MUA rate.

### Position linearization

2D position estimates from the tracking of a head-mounted LED array were linearized by defining a 2D graph representation of the track. We then assigned positions to the most likely edge of the 2D graph using a hidden Markov model (Denovellis et al., 2020). Positions in the central 2D area of the maze were collapsed horizontally onto an axis that ran from the home port, between the two center ports, and ended at the edge of the arms. Nodes designating the start and end of each maze segment (nine segments: the box and eight arms) were manually determined per animal based on the complete position tracking dataset for that subject. Linearized position was binned into 5 cm bins within each segment.

### Spatial decoding

We used a state space model to simultaneously decode the “mental” spatial position of the animal, and whether the position was consistent with a spatially continuous or discontinuous movement model, from unsorted “clusterless” spiking data (Deng et al., 2015; Denovellis et al., 2019; Denovellis et al., 2020; Kloosterman et al., 2014). To do this, we define two latent variables: first, *x*_*k*_, a continuous latent variable that corresponds to the position represented by the population of CA1 cells at time *t*_*k*_ and second, *I*_*k*_, a discrete latent variable that is an indicator for the two movement states we wish to compare: spatially continuous and spatially discontinuous. Our model estimates 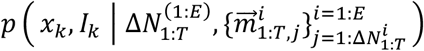, the posterior probability of position and movement state at each time *t*_*k*_ given all the observed multiunit spikes 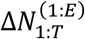 and their associated spike waveform features 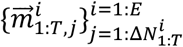 from all CA1 cell layer tetrodes 1: *E*, for all time during a behavioral session 1:*T*. We detected multiunit spikes that had a spike amplitude > 100 μV on at least one tetrode channel. For waveform features, we used the amplitudes on each of the 4 tetrode channels at the time of the maximum detected spike amplitude. Occasionally, our data included tetrodes with 1-2 dead channels; we were able to use these tetrodes by replacing mark amplitudes on dead channels with zeros.

As described in Denovellis et al. 2020, to estimate the posterior probability, we apply a recursive causal filter, starting with initial conditions *p*(*x*_0_, *I*_0_) and iterating from time 1 to time *T*:

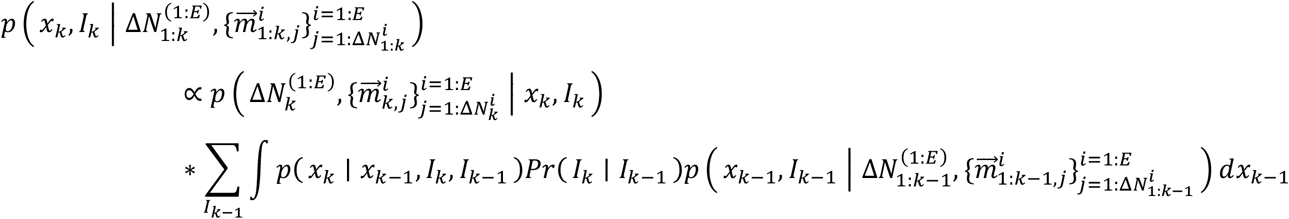

and then apply an acausal smoother, starting from the last estimate of the casual filter 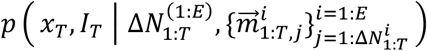 and recursively iterating back to time 1:

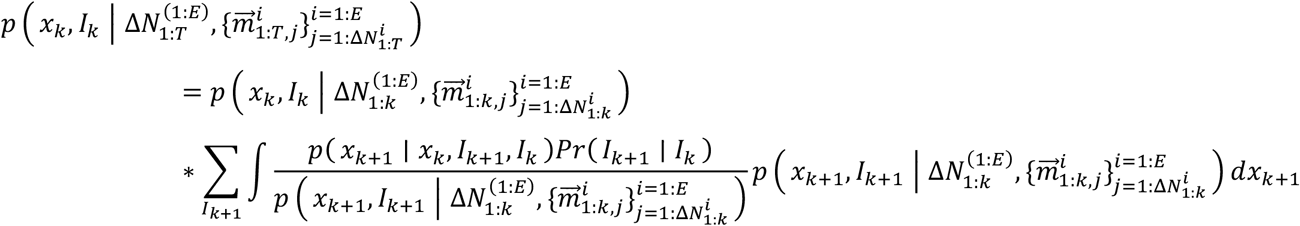

where:

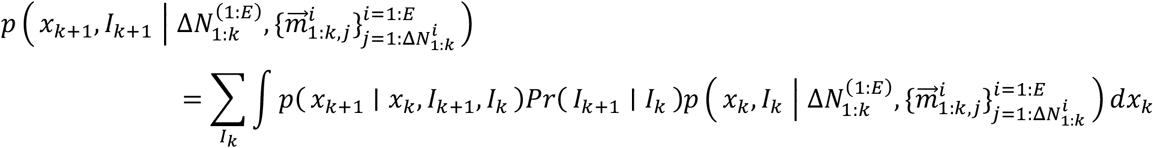

This algorithm requires us to define or estimate four terms: the initial conditions *p*(*x*_0_, *I*_0_), the movement models *p*(*x*_*k*_ | *x*_*k* −1_, *I*_*k*_, *I*_*k* −1_) that differentiate our two states, the discrete transition matrix *Pr*(*I*_*k*_ | *I*_*k* −1_) which defines how likely the model is to switch movement states, and the likelihood of the spiking data 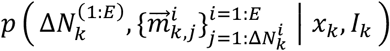. For the initial conditions *p*(*x*_0_, *I*_0_), we set each state *I*_0_ to have equal probability and each initial latent position to have uniform probability density over all possible positions *𝒰*(min *x*, max *x*), reflecting the fact that we do not have any prior knowledge about which state or position is more likely. The two movement states that we include describe two possible ways the latent position could evolve. The spatially continuous movement state predicts that position is equally likely to remain constant or move to a neighboring position bin (spatially continuous) and is approximately defined by a tridiagonal matrix, modified at segment ends. The other state assumes that transitions between all position bins on the maze are equally likely (spatially discontinuous) and is described by a uniform matrix across all position bins (Denovellis et al., 2019). Additionally, we specify that between time steps (2 ms), transitions between the continuous states should reflect the continuous state transition matrix, while transitions between the two states and between discontinuous states should be uniform across all positions. The discrete transition matrix *Pr*(*I*_*k*_ | *I*_*k* −1_) defines how likely the movement state is to switch to the other state versus persist in the same state; we set the probability of persisting in a state to 0.98 and the probability of switching states to 0.02 (Denovellis et al., 2020).

Finally, we specify the likelihood of the observed spikes and associated waveform features (Deng et al., 2016; Kloosterman et al., 2014), which we can compute as:

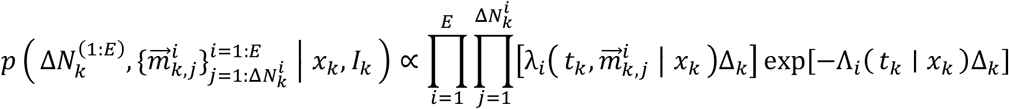

where 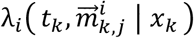 reflects a generalized firing rate that depends on associated waveform features 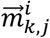 and Λ_*i*_(*t*_*k*_ | *x*_*k*_) is a marginal firing rate that is equivalent to a place field estimated on multiunit spikes. Both of these rates can be defined as:

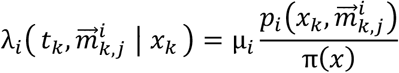

and

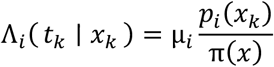

where μ_*i*_ is the mean firing rate for tetrode *i*, π(*x*) is the spatial occupancy of the animal on the track, *p*_*i*_(*x*_*k*_) is the probability of observing a spike on tetrode *i* at position *x*, and 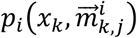 is the probability of observing waveform features 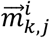 at position *x*. These can all be estimated by training a kernel density estimator on each tetrode’s spike waveform features and corresponding position during the encoding period. The kernel density estimators used a 20 μV bandwidth Gaussian kernel for spike waveform features and 5 cm square bins for position.

We decoded each behavior session independently. For the encoding model, we used all spikes recorded during that session while the subject was moving >4 cm/s. We decoded spiking activity for all times during the behavioral session. When decoding spikes during movement times, the actual position associated with each spike was not used for the decoding of that spike. The resultant joint posterior for each session was a 3D matrix: ∼140 linear positions (5 cm bins) x many 2 ms time steps x 2 movement states. Summed over states, the joint posterior gives a prediction of position for each time bin, as displayed in all example decodes (Figure 2 and Supplementary Figure 3). Alternatively, summed over position, the joint posterior reflects the probability of each movement state over time (Supplementary Figure 3).

Decode quality was assessed by two measures during times of movement (>4 cm/s). First, we computed the fraction of video frames (30 Hz) in which the real position was either on the same maze segment as the prediction of position (MAP; maximum a posteriori), or the adjacent segment (one position was in the box area and the other was in an arm). At these times, which we refer to as “same/adjacent”, the real and predicted positions can be compared unambiguously (in contrast, it is ambiguous whether the distance between locations on two different arms should be measured in linear or 2D distance). The fraction of same/adjacent frames was very high, indicating that the estimated position was usually on the same segment or the adjacent segment as the real position (Supplementary Figure 2A). Second, for same/adjacent frames, we calculated the distance between the predicted and real position (in 5 cm bins; Supplementary Figures 2B and 2C). We excluded two sessions (sessions 1 and 3 of subject 1) due to decoding issues: for the first, a much lower fraction of samples was on the same segment (<75%), and for the second, not all arms were visited during the session, so not all positions were decodable.

We categorized SWRs as spatially continuous or discontinuous based on the probability of each movement state during the event. An event was spatially continuous if the probability of continuous state exceeded a 0.8 threshold (Denovellis et al., 2020) at any point during the event and the discontinuous state did not. In the cases when both states crossed the 0.8 threshold during an event, the event was considered spatially continuous if more time bins were described by the continuous state than by the discontinuous state (for example, see Supplementary Figure 3, SWR 839). Next, to assess which maze segment was most represented during each SWR, we summed the joint posterior across states, calculated the mean posterior over time bins during the SWR, and then summed the posterior density in each maze segment. We considered each spatially continuous event a replay of the segment with the maximum posterior density. Spatially continuous events were further required to have at least 30% mean posterior density in a single maze segment (e.g., an event with 20% density in each of five maze segments would not be considered for further analysis). For all analyses shown, spatially continuous events were categorized as local if the segment with the most posterior density was the same segment as the current location, and remote if it was not. We also repeated all analyses with a more permissive definition, which allowed events that had majority density in the local maze segment to be classified as remote if they had at least 30% posterior density in a remote segment (e.g., an event at the home port with 60% posterior density in the box segment and 40% density in arm 1 would be classified as a remote replay of arm 1). Use of this alternative definition did not alter any of our conclusions.

### Quantification of replay content

To quantify replay of the future, past, previous goal, and current goal arms, we excluded trials during the first trial block of each session, since previous goal was not a valid category during those trials. Within each session, we identified trials of interest: search trials in which past, future, and previous goal were mutually exclusive, and repeat trials in which past, future, and current goal were the same arm, and different from the previous goal arm. Sessions were only included if they contained at least five trials that met these criteria. For each arm category, we counted remote replay events corresponding to the arm category over the trials of interest and divided each count by the total number of remote events during these trials. To calculate the “other arm” fraction, we counted the number of replays corresponding to all “other” arms, divided by the number of possible other arms (for search: five arms; for repeat: six arms), and divided by the total number of remote events. The random distribution was generated by drawing a random integer 1-8 for each remote replay event, counting how often the observed replay corresponded to the randomly drawn arm, repeating this process 5000 times, and then calculating a fraction of total events for each iteration. As expected, this distribution was centered at 0.125 (1/8) with the spread reflecting the number of remote events per subject. For each subject, significance was calculated using Wilcoxon rank-sum test comparing the real fractions of each arm category to the random distribution. The same method was used to quantify replay content for the outer arm events with the exception that local events were included in the analysis of outer arm events. Thus, instead of the fraction denominator being the number of remote events, the denominator was all spatially continuous replay events.

### Generalized linear models to measure the effect of arm category on replay likelihood

Poisson GLMs were fit using the *fitglm* Matlab function. All search and repeat trials after the first trial block of each session were included. One binary predictor was constructed per behavioral category. For instance, for the future arm predictor: each trial contributes eight entries (one per arm), with zeros denoting arms that were not the future arm for that trial and one denoting the arm that was the future arm on that trial. The eight response variables for that trial were the number of times that each arm was replayed on that trial. The model used a log link function, such that log(*replaycount*_*i*_) = *β*_0_ + *β*_1_*x*_1*i*_ + *β*_2_*x*_2*i*_+*β*_3_*x*_3*i*_ where *replaycount*_*i*_ represented the number of times an arm was replayed on a given trial and *β*_0_, the intercept, reflected the baseline replay rate for the subject. The coefficients *β*_1_, *β*_2_, and *β*_3_ represented the weights by which each of the predictor variables *x*_1*i*_, *x*_2*i*_, and *x*_3*i*_ modulated replay rate. For analysis of search trials, the three predictor variables were binary indicator functions denoting whether or not an arm served as the future arm, past arm, or previous goal arm. For analysis of repeat trials, an additional term *β*_4_*x*_4*i*_ was added to measure the effect of the current goal arm. We calculated 99% confidence intervals for the resultant coefficients using Matlab’s *coefCI* and converted them into fold change by exponentiation. Due to the structure of the task, particularly during the repeat phase, it was important to verify that the coefficients were not overly correlated. None of the coefficients were over 50% correlated; correlation matrices for the main search and repeat GLM analyses are reported in Supplementary Figure 4.

### Generalized linear models to predict correct and incorrect repeat phase trials

During the repeat phase, correct trials are those in which the subject chooses to visit the goal arm after having received one or more rewards at the goal location during the current trial block. The trial at the end of each block when the goal has just changed and the subject visits the arm that was the goal arm, but does not receive reward, are considered correct trials. Error trials are those in which the subject visits any arm other than the goal arm after having received one or more rewards at the goal location during the current trial block. For each subject, trials were pooled across sessions and correct trials were randomly subsampled to match the number of error trials. For each subsampled set of correct and error trials, the data was split into five folds; for each fold, one set was held out as a test set and the other four were used as the training set. Binomial GLMs were fit on the training set to classify trials as correct or error (1 or 0) based on either the number of future replay events, the number of current goal replay events, or the numbers of each of the four arm categories: future arm, past arm, current goal arm, and previous goal arm. Misclassification rate was calculated as the mean fraction of the test set that the model classified incorrectly over the five folds. 1000 subsets of trials were drawn from each subject such that the mean and confidence intervals plotted reflect the variability of the mean cross-validated misclassification rate over the 1000 iterations. Because we quantified misclassification rate, values further below 50% reflect better performance.

### Trial-wise replay rate

To quantify replay rate of a particular arm across the transition from rewarded to unrewarded, we first quantified the number of replays per arm across all trials in a session and smoothed the trace with a Gaussian kernel spanning five trials with a standard deviation of one trial. Next, for each transition, we stored the replay rate trace for the goal arm in the window of 40 trials before and after the final rewarded trial of the block. We also stored the replay trace for all arms that were not rewarded during that behavioral session for the same trial window. Trials within the 40-trial analysis window that extended before the start of the session or after the end of the session were simply filled with nans. For the very rare sessions which included more than eight trial blocks, trial blocks past block eight were excluded from analysis, because this would potentially include the goal arm serving as goal twice in the analysis window. Goal arm traces and unrewarded arm traces were compared by performing a Wilcoxon rank-sum test for each trial in the window. Resultant p values were considered significant only after Bonferroni correction for multiple comparisons. The same process was used to compute visit curves, with the exception that the visit trace across trials was not smoothed. The replay rate quantification by search/repeat phase of each trial block was also performed without smoothing of the replay rate trace.

