## Supplemental Data for "Hippocampal replay reflects specific past experiences rather than a plan for subsequent choice"

### SUPPLEMENTARY DATA

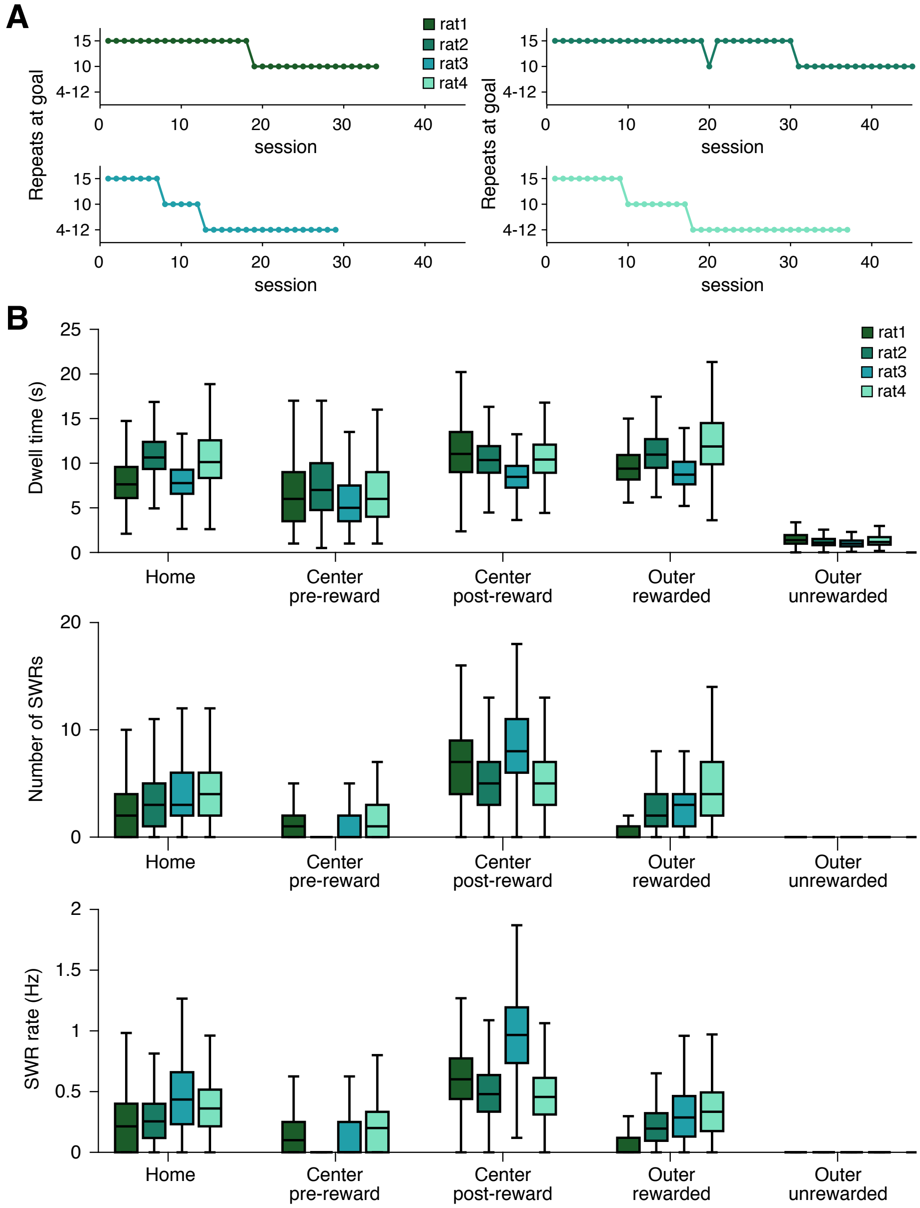


**Supplementary Figure 1. Further characterization of the behavioral task.**
(A) The number of rewards provided at goal arms before the goal would change, for each subject across behavioral sessions. The 4-12 condition indicates that for each session, the reward number was chosen randomly between 4 and 12; all trial blocks within that session would require the same reward number.
(B) For each trial stage (home port, center ports pre- and post- reward, and outer ports on rewarded and unrewarded trials), quantification of the time spent at each trial phase (top), the number of SWRs detected during that time (middle), and the rate of SWRs (bottom). Boxplot range indicates variability in mean values over behavioral sessions; n=24, 32, 23, and 33 sessions per subject, respectively.

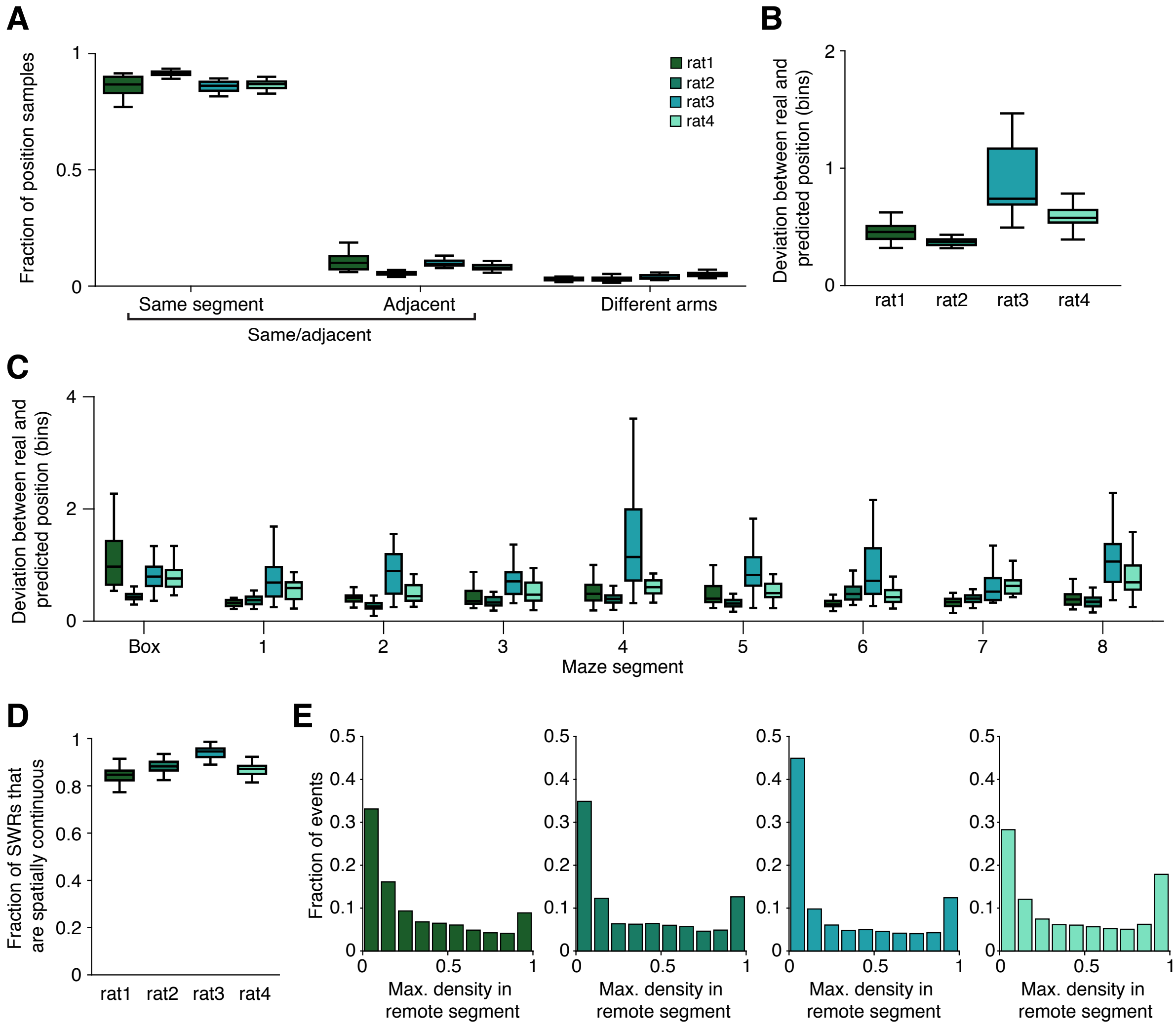


**Supplementary Figure 2. Decoding error during movement times.**

(A) The fraction of samples (30 Hz video frames) during movement in which the actual animal position and the maximum posterior for the decoded position are located in the same maze segment (left), an adjacent segment (one is in the box area and one is in an arm; center), or in two different arms (right). For the first two categories ("same/adjacent"), the distance between the real and predicted positions can be calculated unambiguously (in contrast, it is ambiguous whether the distance between positions on two different arms should be measured in linear or 2D distance).

(B) For each same/adjacent sample during movement, the distance between actual position and the maximum of the posterior is calculated; both are binned in 5 cm increments. Deviation of zero indicates that the real position and the maximum posterior corresponded to the same spatial bin.

(C) For same/adjacent samples during movement, the deviation between real position and posterior separated by maze segment. Deviation is low across all maze segments, indicating that all spatial positions are able to be decoded.

(D) The fraction of SWRs which are considered spatially continuous. Spatially continuous events are SWRs in which more time during the event is spent in the continuous state than in the fragmented state and at least 30% of posterior density is in a single maze segment. Boxplot range indicates variability across sessions; n = 24, 32, 23, and 33 sessions per subject, respectively.

(E) Distribution of the maximum mean posterior density in a single remote maze segment across all events for each subject; n = 22837, 25020, 28289, and 38574 events per subject, respectively.

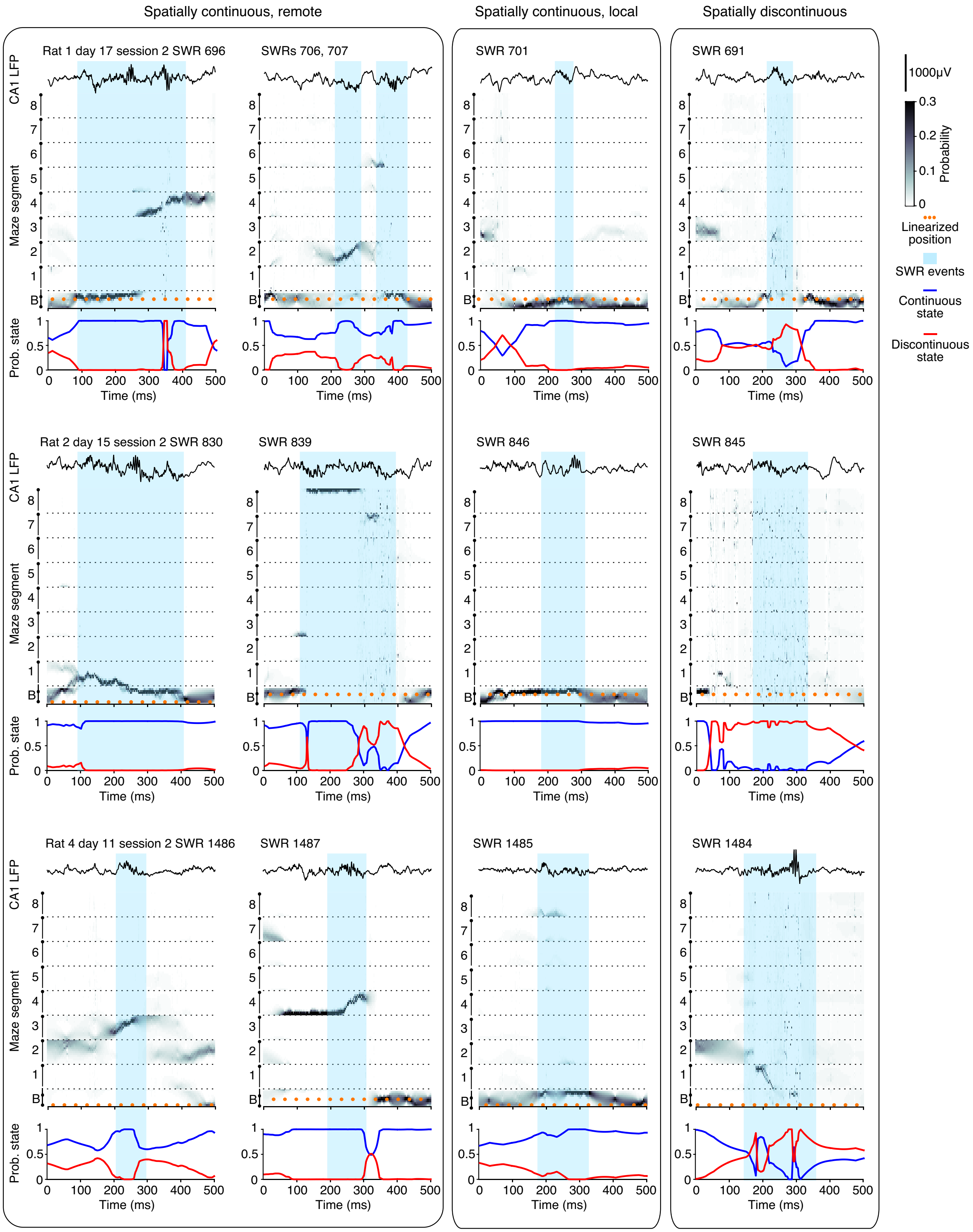


**Supplementary Figure 3. Event examples from additional subjects.**
Each row corresponds to a different subject, with day, behavioral session, and event number indicated. For each subject, the events shown are all drawn from a single trial. The first two columns include spatially continuous remote events. The third column includes spatially continuous local events. The final column includes spatially discontinuous events, which have a majority of time spent in the discontinuous state. Each panel includes a single CA1 LFP trace (top), the posterior and actual position (middle), and finally, the estimated probability of the continuous and discontinuous state variables (bottom) for a 500 ms window surrounding the event. SWR boundaries are indicated by blue shading, and the subject's current position is represented by orange dots.

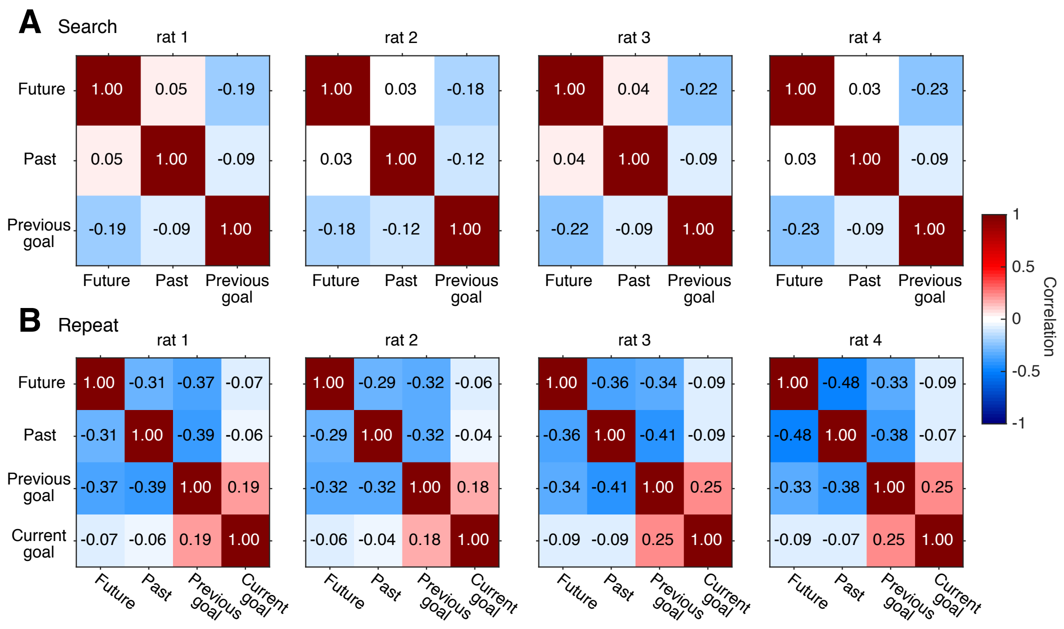


**Supplementary Figure 4. GLM parameter correlation matrices**.
(A) GLM parameter correlation matrices for all search trials, one plot per subject, comparing the coefficients of the three predictor categories.
(B) GLM parameter correlation matrices for all repeat trials, one per subject, comparing the coefficients of the four predictor categories.

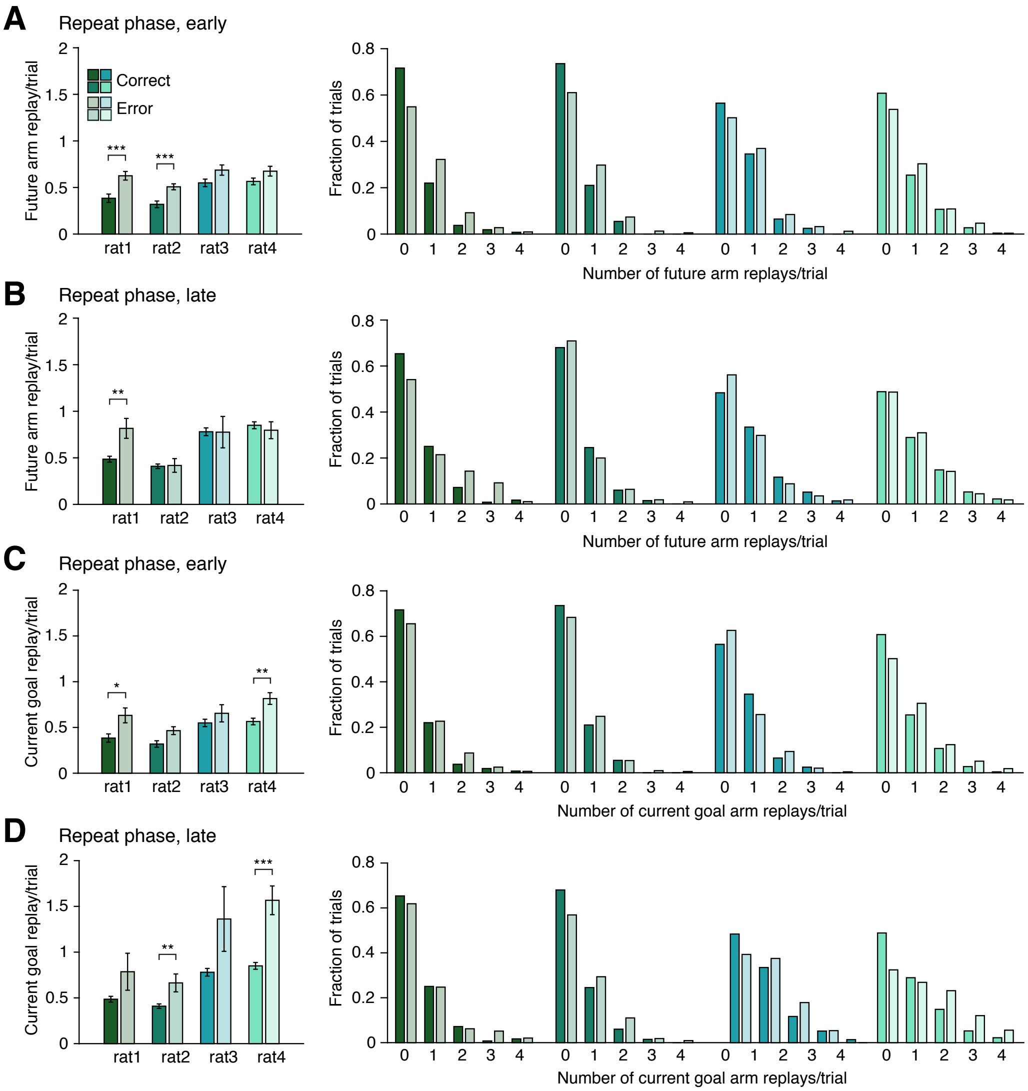


**Supplementary Figure 5. Future and current goal replay on correct and error trials.**
(A) Left, mean rate of future replay per trial during early repeat phase trials (<5 rewards received at the goal location during the current trial block) on correct (solid) and error (shaded) trials. Right, distributions of the number of future replay events per trials on correct and error trials.
(B) Same as (A), but during late repeat phase trials (5 or more rewards received at the goal location during the current trial block).
(C) Left, mean rate of current goal replay per trial during early repeat phase trials (<5 rewards received at the goal location during the current trial block) on correct (solid) and error (shaded) trials. Right, distributions of the number of future replay events per trials on correct and error trials.
(D) Same as (C), but during late repeat phase trials.
For (A) and (C), n = 268, 257, 324, and 515 early correct trials; 326, 544, 249, and 277 early error trials per subject, respectively.
For (B) and (D), n = 655, 750, 523, and 825 late correct trials; 98, 110, 58, and 113 late error trials per subject, respectively.
* denotes p<0.05; ** denotes p<0.01; *** denotes p<0.001, using Wilcoxon rank-sum test. Error bars represent S.E.M.
